# Patterns of speciation are similar across mountainous and lowland regions for a Neotropical plant radiation (Costaceae: *Costus*)

**DOI:** 10.1101/2020.05.30.125757

**Authors:** Oscar M. Vargas, Brittany Goldston, Dena L. Grossenbacher, Kathleen M. Kay

**Affiliations:** Department of Ecology and Evolutionary Biology, University of California, Santa Cruz, Santa Cruz, California 95060; Department of Biological Sciences, Humboldt State University, Arcata, California 95521; Department of Biology, California Polytechnic State University, San Luis Obispo, California 93401

**Keywords:** Ecological specialization, diversification, geographic isolation, Neotropics, pollination shifts, spiral gingers

## Abstract

High species richness and endemism in tropical mountains are recognized as major contributors to the latitudinal diversity gradient. The processes underlying mountain speciation, however, are largely untested. The prevalence of steep ecogeographic gradients and the geographic isolation of populations by topographic features are predicted to promote speciation in mountains. We evaluate these processes in a species-rich Neotropical genus of understory herbs that range from the lowlands to montane forests and have higher species richness in topographically complex regions. We ask whether climatic niche divergence, geographic isolation, and pollination shifts differ between mountain-influenced and lowland Amazonian sister pairs inferred from a 756-gene phylogeny. Neotropical *Costus* ancestors diverged in Central America during a period of mountain formation in the last 3 My with later colonization of Amazonia. Although climatic divergence, geographic isolation, and pollination shifts are prevalent in general, these factors don’t differ between mountain-influenced and Amazonian sister pairs. Despite higher climatic niche and species diversity in the mountains, speciation modes in *Costus* appear similar across regions. Thus, greater species richness in tropical mountains may reflect differences in colonization history, diversification rates, or the prevalence of rapidly evolving plant life forms, rather than differences in speciation mode.

## Introduction

Tropical mountains exhibit extreme species richness and endemism, contribute substantially to latitudinal diversity gradients, and are thought to be cradles of recent speciation (Rahbek et al. 2019a, Rahbek et al. 2019b). The Neotropics contain some of the world’s most species-rich plant diversity hotspots (Barthlott 2005), which all contain substantial mountain ranges. Mountains are hypothesized to play two major roles in the process of speciation: the generation of steep environmental gradients over geographic space (ecogeographic gradients) sensu Gentry (1982) and the geographic isolation of populations by topographic features sensu Janzen (1967). Although studies have linked the timing of montane diversifications with mountain building (Luebert and Wigend 2014), mechanisms by which tropical mountains may promote speciation remain unclear, in part because well-resolved species-level phylogenies for tropical clades remain rare.

Steep montane gradients, in factors such as climate or biotic communities, could promote speciation by ecogeographic divergence without sustained allopatry (Gentry 1982; Angert and Schemske 2005; Hughes and Atchison 2015; Pyron et al. 2015). For example, a marginal population may adapt to novel climate conditions at a species’ upper or lower elevation range limit, at the cost of adaptation to climatic conditions in the remainder of the species’ range (Angert et al. 2008). Similarly, biotic communities (Dobzhansky 1950), such as pollinator assemblages, turnover rapidly in Neotropical mountains (Stiles 1981) and likely contribute to pollinator isolation in plants (Gentry 1982; Kay et al. 2005; Lagomarsino et al. 2016). Taken together, mountains provide large climatic and biotic niche space across short geographic distances, providing an arena for divergent selection and speciation.

Topographic features also can drive allopatric speciation by serving as dispersal barriers regardless of ecogeographic divergence. For example, a species’ range may be divided by a newly formed topographic barrier or individuals may disperse across ridges or valleys to distant areas of suitable habitat. If tropical organisms have narrower climatic tolerances than temperate ones, as hypothesized, the effect of topographic features on isolation may be greatly magnified in tropical mountains (Janzen 1967; Ghalambor et al. 2006; Cadena et al. 2012; Guarnizo and Canatella 2013). Topographic dispersal barriers may lead to frequent progenitor-derivative, or budding speciation, in mountain-influenced areas. This mode of speciation (hereafter, budding) occurs when an initially small colonizing population becomes reproductively isolated from a larger-ranged species (Mayr 1954), and is in contrast to vicariant speciation where a geographic barrier bisects a species’ range (Mayr 1982). Whereas budding speciation may be common in mountains, it is likely less common in lowlands due to fewer steep climatic gradients and topographic barriers.

These long-standing hypotheses about dispersal barriers and ecogeographic gradients in tropical mountains predict unique signatures of speciation. Moreover, if mountains *per se* are driving speciation, patterns of speciation in mountains should differ from lowland regions, which have shallow climatic gradients and less turnover in biotic communities relative to mountains (De Cáceres et al. 2012; Pomara et al. 2012; Fig. 1). First, if ecogeographic divergence is of primary importance, sister species occurring in or around mountains (hereafter, mountain-influenced) are predicted to show climatic niche differentiation (Fig. 1E), and this differentiation should be greater on average than in lowland species pairs (Fig. 1A-B). Similarly, pollinator shifts are predicted to be more frequent in mountain-influenced sister pairs than in the lowlands because of ecogeographic gradients in pollinator assemblages. Second, if topographic dispersal barriers are of primary importance, mountain-influenced sister pairs should frequently show geographic isolation (Fig. 1D), and this isolation should be greater than in lowland species pairs (Fig. 1A), for which geographic isolation may be more ephemeral. If topographic dispersal barriers promote budding speciation, we further predict: 1) greater range size asymmetry between mountain-influenced relative to lowland sister pairs, especially in younger pairs (Barraclough and Vogler 2000; Fitzpatrick and Turelli 2006; Grossenbacher et al. 2014), and 2) nested phylogenetic relationships between recently-diverged mountainous sister species, indicating that small-ranged taxa are derived from widespread progenitors (e.g., Baldwin 2005).

**FIGURE 1.**
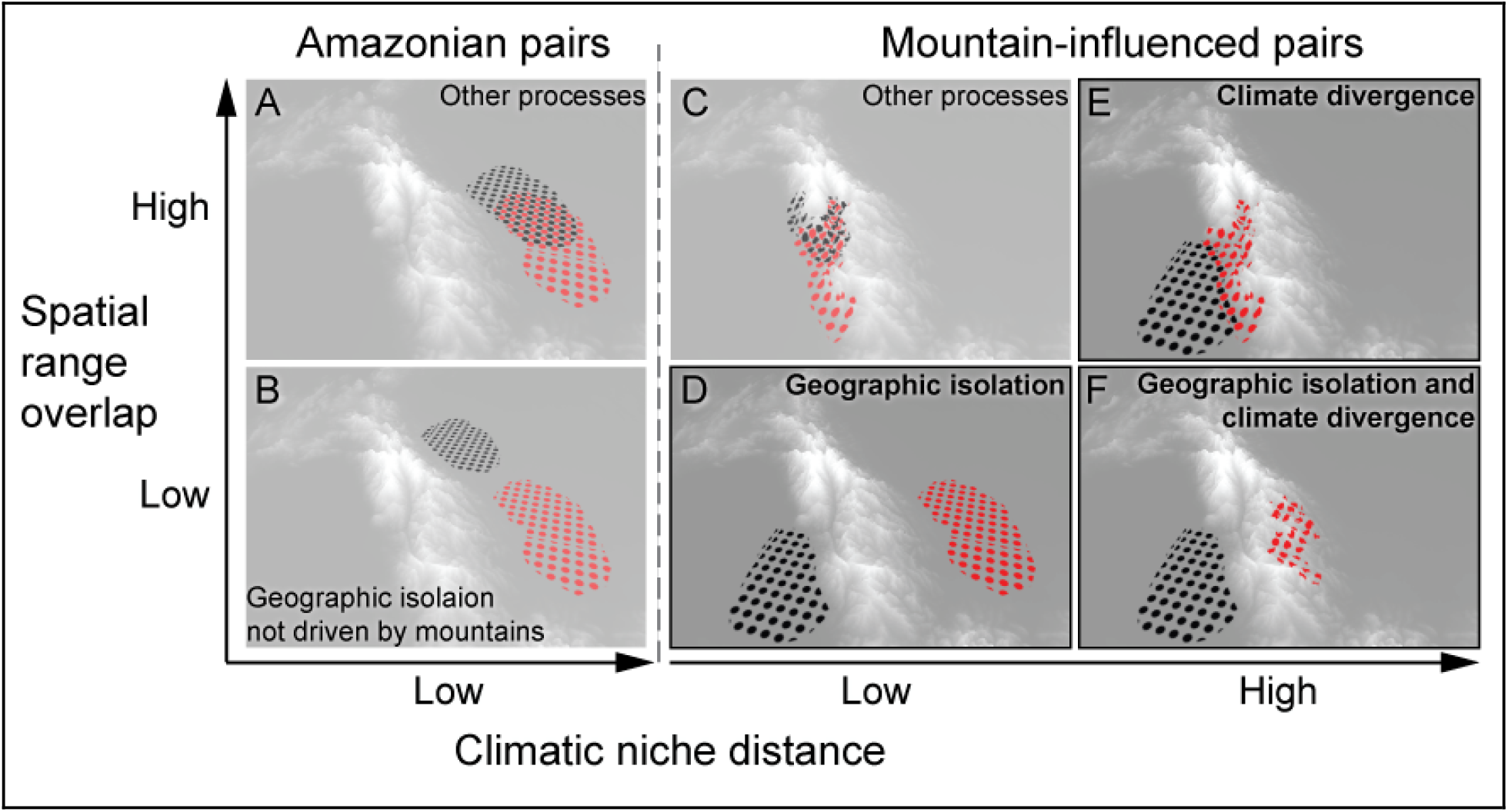
Testing the role of mountains in speciation. Hypothetical ranges of sister species, as black and red filled circles, overlaid on a landscape where lighter grays indicate higher elevations. Sister species in Amazonia, a region with low topographic complexity and relatively homogeneous climate, are expected to be partially sympatric (A) or geographically isolated (B) by lowland geological features, fine-scale habitat divergence, or biotic interactions. Mountain-influenced sister species may also show these patterns (C), however, we predict that geographic isolation (D), climate niche divergence (E), or both (F) will be more common in mountains than in the lowlands (indicated by bolded boxes).

Ecogeographic gradients and topographic dispersal barriers in mountains also predict different patterns of range overlap with divergence time. If allopatric speciation is dominant, then more recently-diverged species pairs should be completely allopatric, whereas older pairs might show range overlap due to range shifts since speciation (Fitzpatrick and Turelli 2006). Contrastingly, if parapatric speciation across ecogeographic gradients is dominant, younger sister species pairs should show partial range overlap whereas older pairs should show a variety of configurations (Fitzpatrick and Turelli 2006; Anacker and Strauss 2014). It is also possible that geographic isolation and niche divergence commonly work together to promote speciation in mountains, with geographically isolated populations adapting to new climate niches (Fig. 1F).

Here we examine speciation modes in the Neotropical spiral gingers (*Costus* L.), a genus comprising approximately 59 species found from sea level to cloud forests throughout tropical Central and South America. *Costus* is a pantropical genus of perennial monocot herbs with a species-rich Neotropical clade nested within the relatively species-poor African taxa. The Neotropical clade likely arose via long-distance dispersal from Africa (Kay et al. 2005, Salzman et al. 2015). Neotropical *Costus* are widely interfertile (Kay and Schemske 2008) with stable ploidy (Maas 1972; 1977). Prior studies have suggested a prominent role for prezygotic reproductive barriers, including ecogeographic isolation (Chen and Schemske 2015), differences in pollination syndrome (orchid bee v. hummingbird; Kay and Schemske 2003), and floral divergence within a pollination syndrome (Kay 2006; Chen 2013). We begin by documenting that *Costus* species richness is indeed higher in Neotropical areas with high topographical complexity, consistent with mountains being strong drivers of speciation in *Costus* (assuming similar extinction rates). We then infer a multi-locus phylogeny for *Costus* that we use to reconstruct the biogeographic history, the timing of divergence, and the evolution of pollination syndromes in the genus. Finally, we use sister species comparisons to test our predictions about speciation modes in montane regions using climatic, geographic, and pollination data. We discuss how our results shed light on speciation in the Neotropics.

## Material and Methods

### OCCURRENCE DATA

We downloaded all known occurrence records for the species in our study from the Global Biodiversity Information Facility (GBIF, http://www.gbif.org). We supplemented these with occurrences from our own field studies in 2018 and 2019, as well as additional herbarium (Cornell, MSU, and UC JEPS) and iNaturalist data (https://www.inaturalist.org). All occurrences were then filtered for quality by excluding records without decimal accuracy in latitude and longitude, and with coordinates failing to match the locality description. To avoid potential taxonomic misidentifications, we retained only occurrences where the identification was made by one of three taxonomic experts: Paul Maas, Dave Skinner, or KMK. We checked species’ epithets against the most recently published taxonomies and corrected synonyms and spelling errors. The final filtered dataset included 4,834 unique occurrences for 61 taxa (mean per taxa=79, range=1-593, SD=121).

### TOPOGRAPHIC COMPLEXITY AND SPECIES RICHNESS

To quantify topographic complexity across our study region, we used the terrain ruggedness index (TRI) function in ArcGIS (ESRI 2018). TRI is a measure of topographic heterogeneity that takes the sum in elevation change between a focal grid cell and all neighboring grid cells (Riley et al. 1999). To calculate TRI, we used raw elevation data at 1km resolution from earthenv.org/topography and projected it to 1600 and 6400 km^2^ grids. The two grid sizes allowed us to assess whether our results were sensitive to spatial scale. Species richness per grid cell was recorded as the number of unique species occurrences in the filtered occurrence data described above (Point Statistics function in ArcGIS). Richness, TRI values, and X,Y coordinates for each grid cell were extracted across the two spatial scales and saved for downstream analysis (Extract Multi Values to Point function in ArcGIS).

To determine whether species richness is predicted by the terrain ruggedness index (TRI), we used both an ordinary least squares regression model and a spatial autoregressive lag model. Ecological data is typically affected by spatial autocorrelation (SAC), with nearby localities being more similar than expected when random (Kissling and Carl 2008). As a result of species distributions being spatially constructed by nature, our data will likely have residual SAC, breaking an assumption of linear regression. To check for residual SAC between TRI and species richness we used Moran’s I coefficient. A Moran’s I coefficient near zero indicates no SAC, while positive and negative values indicate positive or negative autocorrelation (Bhattarai et al. 2004). Spatial autoregressive models are commonly used to mitigate known SAC in the residuals. Accordingly, we used a spatial autoregressive lag model to relate species richness and TRI. Due to the ongoing debate of incorporating spatial autocorrelation into the analysis of species distribution data, we chose to compare our model results to an ordinary least squares regression model which ignores spatial autocorrelation (Dormann 2007). Model selection was accomplished using the Akaike information criterion (AIC).

Because comparisons of species richness may be impacted by uneven sampling across regions (Gotelli and Colwell, 2001), we performed a rarefaction analysis to ensure that our results were not driven by greater sampling effort in mountains relative to lowlands. To do this, we classified all occurrences as either mountainous (TRI>5, N=3477) or lowland (TRI<=5, N=1140). We then randomly drew an equal number of occurrences from the two samples (N=1000), determined richness, and repeated this 1000 times (for a similar approach, see Grytnes and Beaman 2006). Richness in the rarefied samples was considered to significantly differ between mountainous and lowland regions if the 95% confidence intervals were non-overlapping.

### PHYLOGENETIC ANALYSIS

To infer a well resolved phylogeny we employed targeted sequencing, capturing 853 genes in 113 samples representing 57 species, including outgroups (Table S1) and, when possible, samples from different geographic locations for widespread species and putative progenitor-derivative species pairs. We selected the genes based on six transcriptomes belonging to neotropical *Costus*, five newly sequenced (Table S2) and one published (GenBank BioSample: SAMN00991785). We extracted total RNA using the RNeasy Plant Mini Kit (Qiagen, San Diego, California) from fresh tissue; Poly-A enriched libraries (with an insert size of 300 bp) were prepared by the DNA Technologies Core at the University of California, Davis, with subsequent paired-end 150 bp HiSeq4000 sequencing (Illumina Inc.). We employed SeqyClean v.1.10.07 (Zhbannikov et al. 2017) to remove low quality reads and read tails using default parameters and a cutoff Phred score of 20. Poly A/T tails were also trimmed. Assembly of transcripts was performed with Trinity v2.8.4 (Grabherr et al. 2011). With the transcriptomic dataset, we employed Captus (https://github.com/edgardomortiz/captus) to select the genes for sequencing. Briefly, Captus first used VSearch V.2.10.3 (Edgar 2010) to deduplicate individual transcriptomes (query_cov [overlap] = 0.99, id [similarity] = 0.995) and performed clustering among the transcripts of all samples (query_cov = 0.75, id = 0.75) outputting a fasta file for every cluster. Clusters were subsequently aligned with MAFFT v7.407 (Katoh and Standley 2013) using the “--auto” mode. Only genes for which a single copy was found in the alignments were used for subsequent subselection, filtering out possible paralogs. Final gene selection for sequencing in the phylogenetic analysis was based on transcript length (len = 720-2400), transcript presence in a minimum of four species (spp= 4,6), transcript presence in a focal species (*Costus pulverulentus*, foc = 1), percentage of gaps in the alignment not exceeding 50% (gap = 50), an average pairwise percentage identity range of 75–99.6% (pid = 75-99.6), and allowing a maximum of 15% of short introns per gene (< 120 bp) (psr = 0.15). Finally, allowing for a G-C content of 30–70% and a tailing percentage overlap of 66.55, Captus designed 16,767 baits of 120bp in length for the 853 genes selected.

We extracted DNA from recently collected field and greenhouse samples using NucleoSpin Plant Mini Kit II (Macherey-Nagel, Düren, Germany) according to the manufacturer’s protocol, adding 5 uL proteinase K (20 mg/mL) to the digestion step and increasing the digestion incubation time to an hour. For herbarium specimens we used the MagPure Plan DNA LQ kit (Angen Biotech, Guangdong, China). Library preparation and sequencing for the 853 targeted genes was performed by Rapid Genomics (Gainesville, Florida). We employed HybPiper v1.3.1 (Johnson et al. 2016) to assemble the targeted genes, and MAFFT using the “linsi” exhaustive algorithm to align the matrices containing concatenated exons and introns. Problematic sections in the alignments were trimmed with the “-automated1” option of trimAl v1.4.rev22 (Capella-Gutiérrez et al. 2009). Rogue taxa were removed with the “-resoverlap 0.75 -seqoverlap 75” arguments of trimAl.

To filter out tentative paralog genes unidentified by Captus, we excluded genes with extreme variation in branch lengths based on the assumption that ingroup branches should not be extremely long considering the recent diversification of Neotropical *Costus* (Kay et al. 2005). To identify genes with extreme in-group branch length variation, we first inferred trees for each alignment with IQ-Tree v1.6.12 (Nguyen et al. 2015) using a GTR+G model and 1000 ultrafast bootstraps. Then, after removing outgroups with pxrmt (Brown et al. 2017) and outlier sequences with TreeShrink “-q 0.10” (Mai and Mirabab 2018), we calculated the variation in branch lengths using SortaDate (Smith et al. 2018), and then sorted genes accordingly. Visual examination of genes with extreme variation in branch lengths revealed possibly paralogy. Therefore, to be conservative we filtered out genes in the top 10% distribution of branch-length variation, resulting in 756 genes for subsequent phylogenetic analysis. Visual examination of remaining genes after filtering revealed no potential paralog issues.

We used concatenated- and coalescent-based approaches for the inference of species trees. Before concatenation, gene alignments were filtered from outlier sequences flagged previously by TreeShrink in our gene trees. A matrix containing sequences for 756 genes was used to infer a concatenated-based species tree, using IQ-Tree with an independent GTR+G model of sequence evolution for each gene partition and 1000 ultrafast bootstraps. We calculated the number of gene trees supporting a given node in the concatenated topology by employing phyparts (Smith et al. 2015), and results were plotted with phypartspiecharts.py (https://github.com/mossmatters/MJPythonNotebooks). For the coalescent inference, we inferred a species tree based on the 756 IQ-Tree-inferred topologies with ASTRAL v.5.6.3 (Zhang et al. 2017), collapsing branches with less than 90 ultrafast-bootstrap support and removing from each input tree the taxa flagged as having outlier branch lengths.

To determine whether *Costus* diversification coincided with substantial mountain uplift, and to estimate divergence time for sister species, we calibrated our concatenated topology using fossils and external *non-Costus* Zingiberales sequences (Table S3). The inclusion of *non-Costus* sequences was necessary because of the absence of *Costus* fossils. First, we identified the top 50 most clock-like genes in our dataset using the metrics outputted by SortaDate, considering in order of priority branch-length variance (low variance preferred), root-to-tip length (high length preferred), and topological similarity with the concatenated topology (high similarity preferred). Then, we mapped filtered transcriptomic reads from nine Zingiberales species and two outgroups to the top 50 most clock-like genes using reads2sam2consensus_baits.py (Vargas et al. 2019), which wraps sam2consensus.py (https://github.com/edgardomortiz/sam2consensus). The resulting matrices with *Costus* and non-*Costus* sequences were filtered for those containing all non-*Costus* taxa and reduced by leaving only one sample per monophyletic species. We aligned the 27 resultant gene matrices with MAFFT using the “linsi” algorithm and filtered the alignments for 95% column occupancy with the command “pxclsq” of Phyx (Brown et al. 2017). The filtered 27 alignments were concatenated and analyzed with BEAST v.2.6.1 (Bouckaert et al. 2014) with independent GTR+G models for each gene partition. As a prior, we used a chronogram with relationships fixed based on a Zingiberales phylo-transcriptomic analysis (Carlsen et al. 2018) and our concatenated tree. Branch lengths for the prior tree were calculated with IQ-Tree and later parameterized with TreePL (Smith and O’Meara 2012). Calibrations points were set as follows: 69 Mya (CI = 63–76) to the stem node of the Zingiberaceae based on the fossil *Zingiberopsis magnifolia* (Hickey and Peterson, 1978), and 77 Mya (CI = 69–86) to the crown clade of the Zingiberales based on *Spirematospermum chandlerae* (Friis 1988). Using a birth-death model, BEAST was set to run for 100 m generations sampling every 1 k. We calculated the chronogram after combining 6 sets of 2500 trees from independent runs with LogCombiner v.2.6.1, inputting those in TreeAnnotator v.2.6.1 (Bouckaert et al. 2014) after checking for chain convergence and a minimum effective sample size of 200 for all parameters with Tracer 1.7.1 (Rambaut et al 2014). We used FigTree v1.4.4 (https://github.com/rambaut/figtree/releases) and ggtree to produce the tree figures (Yu et al. 2017).

### BIOGEOGRAPHIC ANALYSIS

To test whether early ancestors of *Costus* originated in the mountains or lowlands, we inferred the biogeographic history of the group. We first scored the absence and presence of extant species in four bioregions, Central America + Choco (C), West Indies (W), Andean (A), and Amazonian (M), based on our curated occurrence dataset. We then performed an ancestral range reconstruction using the dispersal-extinction-cladogenesis model (DEC; Ree and Smith 2008), a likelihood version (DIVALIKE) of the dispersal-vicariance model (Ronquist 1997), and a likelihood implementation (BAYAREALIKE) of the BAYAREA model (Landis 2013) as implemented in BIOGEOBEARS (Matzke 2013). We abstained from using the founder J parameter given its caveats (Ree and Sanmartín 2018). BIOGEOBEARS infers ancestral areas using the aforementioned models and compares them based on likelihoods. Bioregions were modified from a previous biogeographic study of the Neotropical region (Morrone 2014), considering the distribution of *Costus*. Our input tree was the chronogram inferred after time calibration analysis. The most likely reconstruction was selected based on the corrected Akaike Information Criterion (AICc).

### ESTIMATING CLIMATE NICHE

To estimate the climate niche of each species, we obtained four variables representing aspects of temperature and precipitation (http://www.worldclim.org/): mean annual temperature, mean annual precipitation, temperature seasonality, and precipitation seasonality. All data were projected into a South America Albers Equal Area Conic projection and resampled to a 1 km x 1 km grid cell size. Realized niche position of each species was estimated by circumscribing each species’ occurrence-based niche relative to all occupied niche space across Neotropical *Costus* using the “PCA-env” ordination technique implemented in the Ecospat package (Broennimann et al. 2012; Broennimann et al. 2018). Here, the dimensions of the environmental space for *Costus* were reduced to the first and second axes from a principal components analysis (PCA). The PCA of the four climate variables was constructed using all curated Neotropical *Costus* occurrences, subsampled to one occurrence per grid cell (N=2743 grid cells total). We then created a grid with 100 x 100 PCA unit grid cells and used the species’ presence data to project the density of each species into environmental space using a kernal density function (Broennimann et al. 2012). Niche position for each species was estimated as the mean of PC1 and PC2.

To determine whether mountain-influenced taxa occupy a larger volume of climate niche space overall, we tested whether the variation in species’ mean niche values differ by region. Species were categorized as either Amazonian (all occurrences contained in the Amazon and/or West Indies bioregions) or mountain-influenced (occurrences fully or partially contained in the Central America + Choco or Andean bioregions). We visualized the evolution of climate niches by projecting the phylogeny on species’ mean values for PC1 and PC2 using Phytools (Revell 2012), and we used Levene’s tests on PC1 and PC2 separately to compare the variance in mean niche values between regions (leveneTest function, car package in R, Fox and Weisberg 2019).

### EVOLUTION OF POLLINATION SYNDROMES

We evaluated whether hummingbird pollination and/or shifts from orchid bee to hummingbird pollination are more prevalent in mountain-influenced than Amazonian taxa. Orchid bee- or hummingbird-pollination syndromes were assigned to taxa based on previous studies in the genus (Maas 1972; Maas 1977; Kay and Schemske 2003; Kay et al. 2005) and KMK expertise. Although only a subset of species has pollination data, pollination syndromes accurately predict whether orchid bees v. hummingbirds are the primary pollinator (Kay and Schemske 2003). Thus, pollination syndromes serve as a tractable proxy for an important biotic interaction that could contribute to ecogeographic divergence. We first used a chi-squared test to determine whether hummingbird pollination is more frequent among mountain-influenced species than Amazonian species, regardless of their phylogenetic history. To account for phylogenetic history, we used Pagel’s (1994) test to determine whether there was correlated evolution of pollination syndrome and geographic region (function fitPagel, phytools package, Revell 2012; although see Maddison and FitzJohn 2015 for caveats regarding this method). We then performed a character reconstruction of pollination syndromes on the phylogeny (function make.simmap, phytools package; Revell 2012) and used another chi-squared test to determine whether reconstructed shifts from bee to hummingbird pollination are more likely in mountain-influenced vs. Amazonian ancestors. Ancestors were categorized based on the biogeographic and pollination character state reconstructions.

### IDENTIFYING SISTER TAXA

We identified sister taxa for comparisons of range overlap, climate niche divergence, and pollination shifts in recent and phylogenetically independent speciation events. Sister taxa were identified from the 1000 rapid bootstraps of the concatenated alignment by first pruning the trees to the reduced taxon set used for the time-calibrated phylogeny and then counting the frequency of all sister pairs across bootstrap replicates with a custom R script (R Core team 2020). This frequency was used as a weighting factor in downstream analyses to account for uncertainty in tree topology. Sister pairs were categorized as either mountain-influenced (one or both species occurred in Central America, Choco, or the Andes) or as Amazonian (both species occurred in the lowland Amazon, or one species in the Amazon and the other in the West Indies). We flagged potential cases of budding speciation in the phylogeny when we observed a widespread species having a taxon nested in it with a smaller range area; we arbitrarily chose a minimum asymmetry ratio of 5 (large / small range) as a cut off.

### ESTIMATING SISTER PAIR RANGE OVERLAP, RANGE ASYMMETRY, AND CLIMATIC NICHE DIVERGENCE

For each sister pair, we used the filtered occurrence data to estimate the degree of range overlap using a grid approach. We divided the Neotropics into a series of cells by grid lines that follow degree longitude and latitude using the “raster” R package version 2.9-5 (Hijmans 2016). We calculated range overlap as the summed area of grid cells occupied by both species, divided by the summed area of occupied grid cells for the smaller ranged species. Thus, range overlap could range between 0 (no range overlap) and 1 (the smaller-ranged species is found only within the range of the larger-ranged species) (Barraclough and Vogler 2000; Fitzpatrick and Turelli, 2006). We calculated range size asymmetry as the summed area of grid cells occupied by the larger ranged species divided by the summed area of grid cells for the smaller ranged species (Fitzpatrick and Turrelli 2006). In order to assess whether the ensuing analyses were sensitive to spatial scale, range overlap and size asymmetry were calculated at two cell sizes, 0.05 and 0.1 decimal degrees, representing grid cells of approximately 33 and 131 km^2^ respectively (exact value varies by latitude). Sister pairs lacking adequate geographic data (fewer than 4 known occurrences for one or both species) and those taxonomically poorly understood were excluded from all downstream analyses (Table S4).

### COMPARING MOUNTAIN-INFLUENCED AND AMAZONIAN SISTER PAIRS

We performed a series of analyses to determine whether climate divergence or geographic isolation differs between mountain-influenced and Amazonian sister pairs. We predicted that mountain-influenced pairs would have greater niche divergence and/or greater geographic isolation (less range overlap), and more frequent budding speciation than Amazonian pairs. We also predicted that mountain-influenced pairs would have more frequent shifts to hummingbird pollination than Amazonian pairs, but were unable to make this comparison with only sister pairs because of the small number of shifts occurring at the tips of the phylogeny.

To compare climate niche divergence between regions, we compared the frequency of climate niche equivalency and the degree of climate niche divergence for mountain-influenced v. Amazonian sister pairs. To estimate climate niche equivalency, we determined whether each sister pair occupies statistically equivalent niches, i.e., that the niche overlap between sister species is equal to that of two species occupying random niches in the same range of environmental conditions that are available to the species in question (Warren et al. 2008; Broennimann et al. 2012). This was performed using the function ecospat.niche.equivalency.test (ecospat package, Broennimann et al. 2018), whereby the observed overlap is compared to a null distribution of simulated overlaps when randomly reallocating the occurrences of both species among the joint distribution of occurrences. Only pairs where each member species occupied at least 5 grid cells were used in this analysis (N=22 sister pairs). The frequency of climate niche equivalency was compared between regions (mountain-influenced, Amazonian) with a weighted chi-squared test, weighted by the number of bootstrapped trees containing a given sister pair (function wtd.chisq, weights package, Pasek 2020). To compare the mean climate niche divergence of sister pairs between regions, we calculated climate niche divergence for each pair as the euclidean distance between mean PC1 and PC2 for each species and then used a two sample T-test, weighted by the number of bootstrapped trees containing a given sister pair (wtd.t.test function, weights package, Pasek 2020).

To compare geographic isolation between regions, we first quantified the current range overlap for sister pairs between regions and then examined how range overlap varies with divergence time. To compare the mean current range overlap, we used a two sample T-test, weighted by the number of bootstrapped trees containing a given sister pair (wtd.t.test function, weights package, Pasek 2020). We note that current overlap may differ from overlap at the time of speciation due to post-speciation range expansions, contractions and shifts; however, by using only sister species and comparing regions, we can infer differences in geographic isolation between regions for the most recently diverged species pairs in *Costus*.

To determine whether allopatric speciation is more prevalent in the mountains, we tested whether sister pair range overlap was predicted by divergence time, region, and their interaction using a linear model (lm function, weighted by the number of bootstrapped trees containing a given sister pair, R). If allopatric speciation is dominant, then more recently-diverged species pairs should be allopatric, whereas older pairs might show range overlap due to range shifts since speciation (Fitzpatrick and Turelli 2006). Contrastingly, if parapatric speciation is dominant, younger sister species pairs should show range overlap whereas older pairs should show a variety of configurations (Fitzpatrick and Turelli 2006; Anacker and Strauss 2014). A significant interaction between divergence time and region would indicate that the predominant geographic mode of speciation differs by region.

Finally, to determine whether budding speciation occurs and whether this phenomenon varies by region, we examined the relationship between range asymmetry and divergence time, and we also looked for evidence of nested phylogenetic relationships that would indicate a small-ranged taxon was derived from within a widespread progenitor taxon. We first tested whether sister pair range asymmetry was predicted by divergence time, region, and their interaction using a generalized linear model with a natural log link function, gamma distribution suitable for left-skewed response variables such as range size asymmetry (glm function, weighted by the number of bootstrapped trees containing a given sister pair). If budding speciation is common, then range size asymmetry is predicted to be greatest for the youngest sister pairs and to decrease on average with time, as ranges undergo expansion or contraction following the initial budding speciation event (Fitzpatrick and Turelli 2006; Grossenbacher et al. 2014). A significant interaction between divergence time and region would indicate that the signature of budding speciation differs by region. Significance of predictors was assessed by likelihood ratio chi-squared tests (LR) using single term deletions. Our phylogenetic sampling also allowed us to assess nested phylogenetic relationships indicative of budding speciation in four putative cases, three mountain-influenced and one Amazonian (*C. scaber* in Central America – *C. ricus, C. pulverulentus – C*. sp. nov. 18020/18049, *C. laevis – C. wilsonii*, and *C. scaber* in South America – *C. spicatus*, respectively). We did not know in advance which, if any, species were produced by budding speciation, but we attempted to sample across the known geographic distributions of multiple widespread taxa.

Because we only have a modest set of sister pairs that are used repeatedly in the analyses above, we present exact p-values and describe effect sizes, without using a strict α = 0.05 to determine significance.

## Results

### Topographic complexity and species richness

We find a positive association between topographic complexity and species richness in *Costus*, consistent with mountains playing a key role in diversification of this clade (Fig. 2). We recover a significant spatial autocorrelation between Terrain Ruggedness Index (TRI) and species richness (Fig. 2; Moran’s I > 0.12, P < 0.001, across both spatial scales). Ordinary least squares regression shows a significant positive relationship between TRI and species richness (1600 km^2^, F = 7.56_1,973df_, P = 0.006; 6400 km^2^, F = 6.82_1,597df_, P = 0.009). Simultaneous Autoregressive (SAR) lag models also shows a positive relationship between TRI and species richness, however, significance varies by scale (1600 km^2^, z=1.89, P = 0.138, pseudo-r squared = 0.227; 6400 km^2^, z = 1.89, P = 0.049, pseudo-r squared=0.246). SAR lag models are favored over ordinary least squares regression using AIC (1600 km2, Likelihood Ratio = 1282.51, P < 0.001; 6400 km2, Likelihood Ratio = 170.55, P < 0.001). Overall, *Costus* shows a center of species richness in the Central America + Choco and the northern Andean floristic regions, moderate richness in the Guiana Shield and the eastern slope of the southern Andes, and relatively low richness in the Amazon basin. It is important to note that our sampling relied on collection efforts carried out by previous collectors and our team, which largely focused on sampling Central America, and therefore it is possible that Andean and Amazonian *Costus* diversity is underestimated. Finally, after accounting for uneven sampling between regions using rarefaction, we find that richness is significantly greater in mountainous than lowland regions (mean rarefied richness = 55.6 and 40.3 species respectively and 95% confidence intervals are non-overlapping, Fig. S1).

**FIGURE 2.**
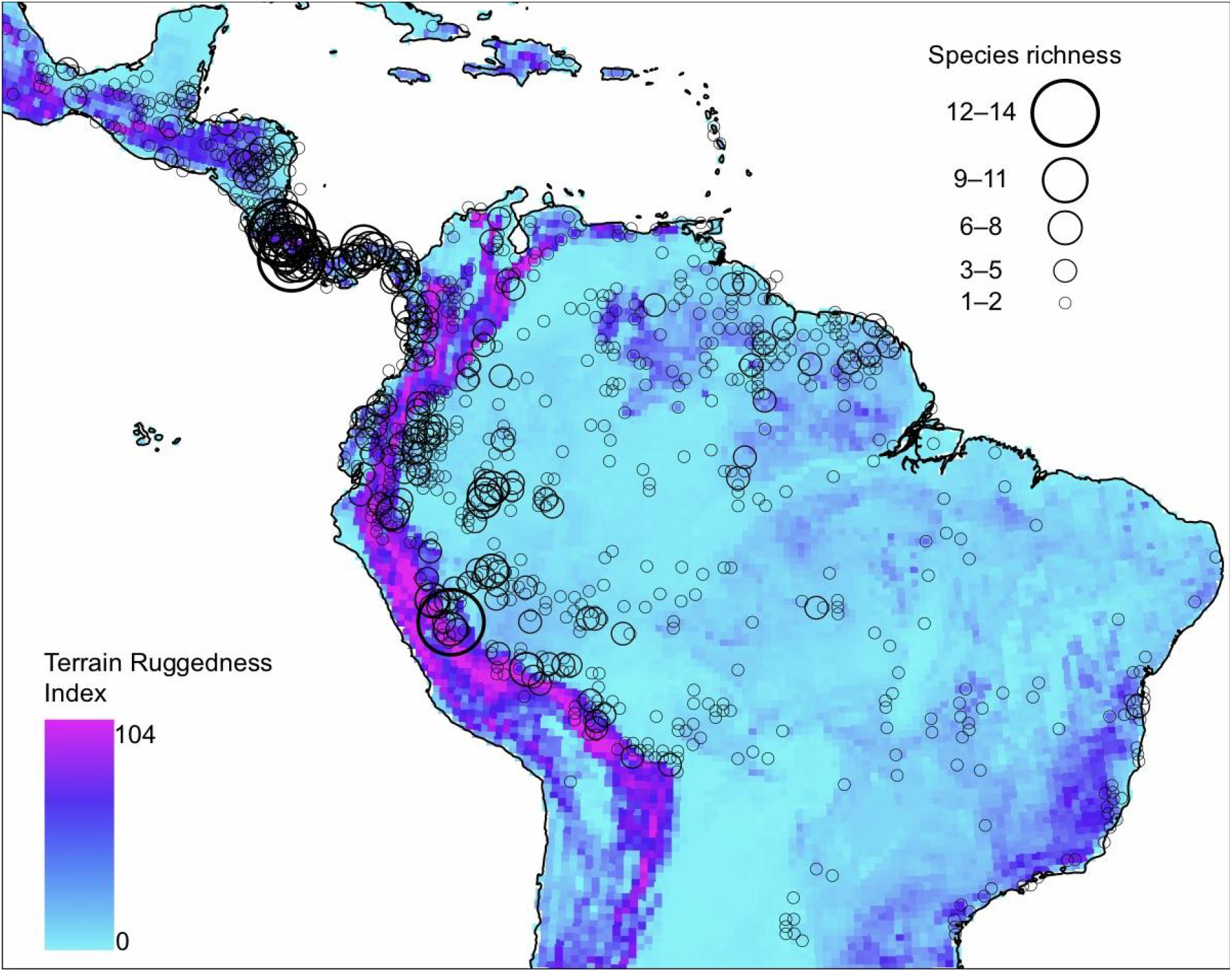
Species richness mapped onto a grid measuring topographic complexity using the Terrain Ruggedness Index. Warmer colors indicate higher topographic complexity, and circles are scaled to the number of reported species per grid.

### Phylogenetic analysis

We infer a robust species-level phylogeny for *Costus*. Our concatenated matrix has 95.3% cell occupancy comprising 133 samples by 756 genes in 1,474,816 aligned columns (Tables S5–6). Average length per gene, including partial introns, is 1,951 bp. Samples have an average of 728 genes. We calculate two species trees, the first based on a concatenated alignment and a second based on individual gene trees in a coalescent framework. Both species trees have robust support and similar topologies (Figs. 3,S2). Because of the general low molecular divergence found in the ingroup (average pairwise identity = 90.0%) and the low signal found in individual genes (Fig. S3), we selected the concatenated topology as the best phylogenetic hypothesis for the remainder of this study (Fig. 3). Our *Costus* phylogeny is robust with most nodes presenting full or high support (>= 95 ultrafast bootstrap, >= 90 Shimodaira-Hasegawa approximate likelihood ratio test) and generally agrees with previous phylogenetic studies in *Costus* (Kay et al 2005, André et al 2016).

**FIGURE 3.**
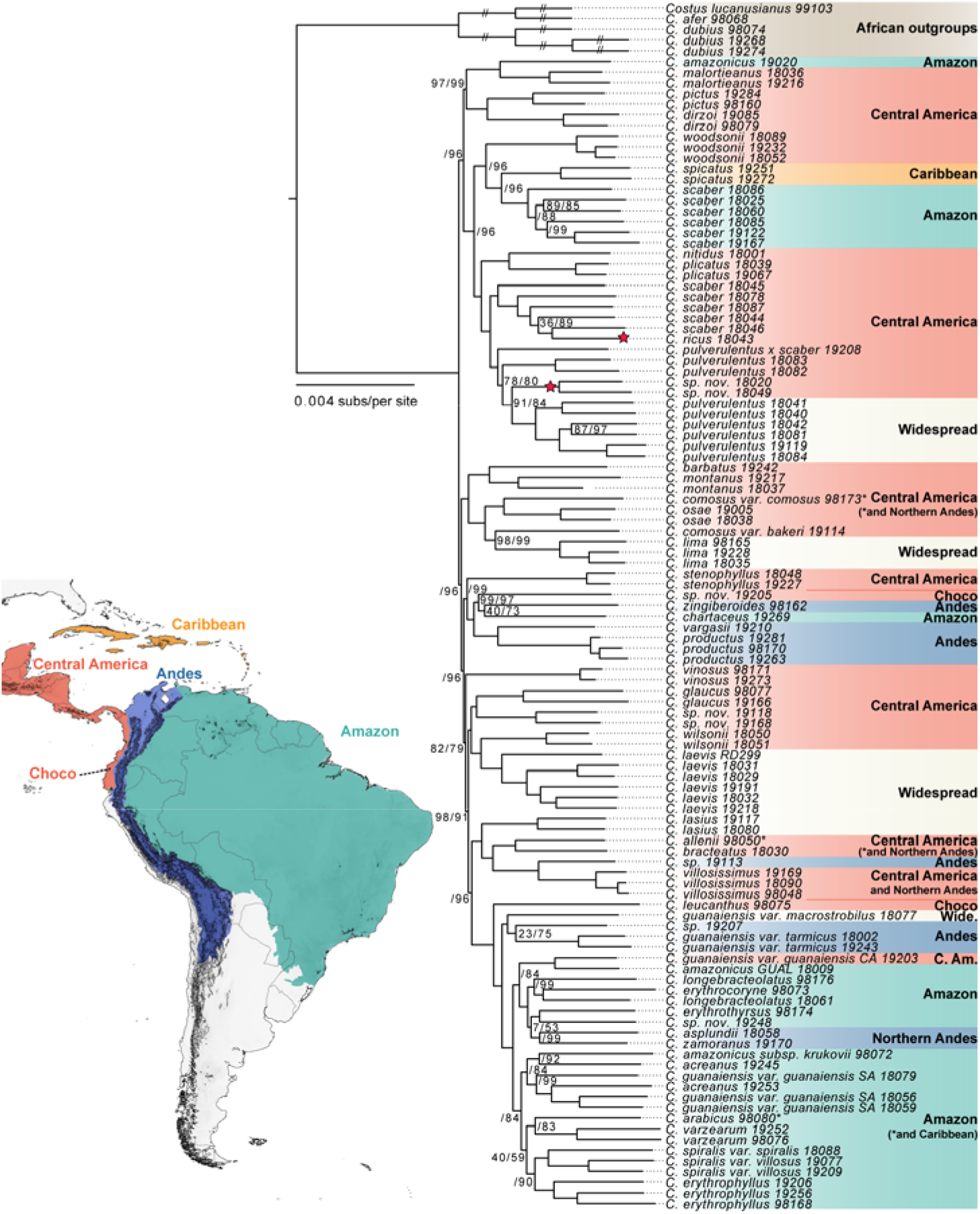
Phylogram inferred from the concatenated matrix of 756 genes for *Costus*. Node numbers indicate ultrafast bootstrap (left) and a Shimodaira-Hasegawa approximate likelihood ratio test (right). nodes without values shown are fully supported at 100. Outgroup branch lengths are reduced to save space. Terminal clades are color coded according to the geographic region on the map. Stars indicate species with evidence for budding speciation.

### TIME-CALIBRATION OF THE PHYLOGENY

Our time-calibrated phylogeny dates the crown clade of Neotropical *Costus* to 3.0 Mya with a 95% CI = 1.50–4.87 (Fig. 4). Our matrix for the calculation of the chronogram, which included Zingiberales sequences for each one of its families and a reduced *Costus* sampling, is composed of 69 taxa by 27 clock-like genes, comprising 22,237 aligned columns (Table S7). Our time-estimation for the origin of *Costus* in the Neotropics is consistent with a previous dating of 1.1–5.4 Mya (Kay et al. 2005) but younger than another of ~7 Mya (André et al. 2016).

**FIGURE 4.**
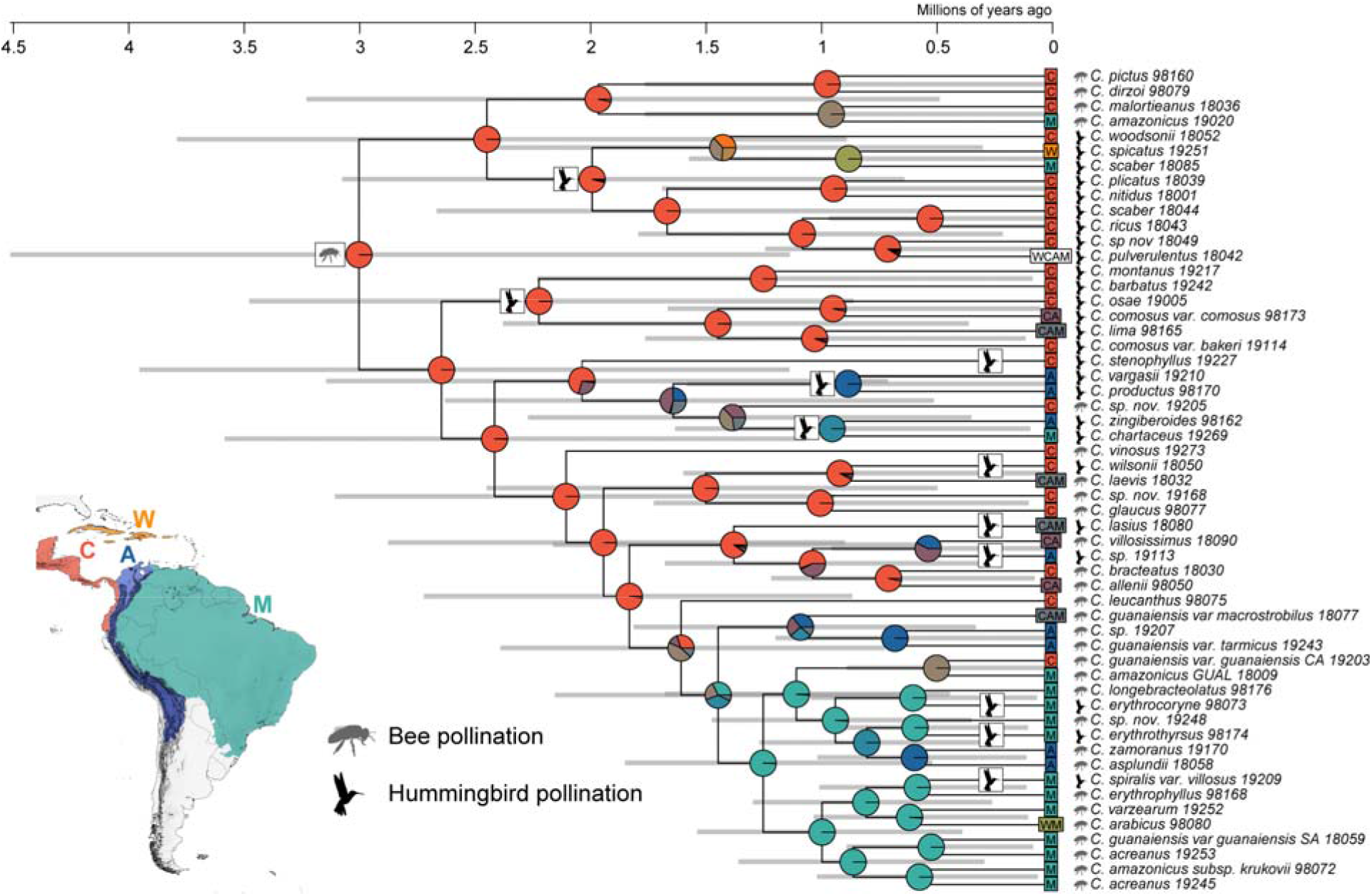
Time calibrated phylogeny, DIVALIKE biogeographic historical inference, and ancestral pollination character reconstruction for Neotropical *Costus*. Pie charts represent percent probabilities of areas for the ranges of ancestors. C: Central America and/or Choco, W: West Indies, A: Andes, M: Amazon. Mixed colors indicate combined bioregions. Pollination syndromes are indicated for every tip taxon along with inferred transitions from bee to hummingbird pollination on internal branches. The most recent common ancestor is reconstructed as bee pollinated. Gray horizontal bars represent 95% confidence intervals for node ages.

### BIOGEOGRAPHIC ANALYSIS

Ancestral range reconstruction suggests that the Central American region has dominated the biogeographical history of the genus in the Neotropics (Fig. 4). Based on AICc, the model that best fits the reconstruction is DIVALIKE (AIC = 185.5), followed by DEC (AIC = 188.2), and BAYAREALIKE (AIC = 230.5). The DIVALIKE reconstruction shows that nearly half of the ancestors in the phylogeny were distributed in Central America (25 out of 54 ancestors with a > 0.90 probability) with the vast majority of the early ancestors estimated as Central American. Colonization out of Central America is inferred to have happened around 1.5 Mya to the Andean and the Amazon regions. Similar biogeographical patterns are also found in the reconstruction inferred with the DEC but not BAYAREALIKE models (Figs. S4,S5).

### CLIMATE NICHE OF MOUNTAIN-INFLUENCED AND AMAZONIA SPECIES

Principal component analysis reveals the first two climate niche axes explain 42 and 23% of the variation among all *Costus* occurrences, respectively. PC1 primarily describes variation in mean annual precipitation and seasonality: low values indicate greater precipitation and high values indicate higher seasonality in both temperature and precipitation. PC2 primarily describes variation in mean annual temperature: low values indicate cooler environments. Overall, we found that the variance in species’ mean niche values was greater among mountain-influenced than among Amazonian species for PC2 (Fig. 5; PC2 F = 10.05^1,46df^, *P* = 0.003), but not PC1 (Levene’s test: PC1 F = 0.05^1,46df^, *P* = 0.824), indicating that, together, mountain influenced taxa are occupying a larger temperature niche space.

**FIGURE. 5.**
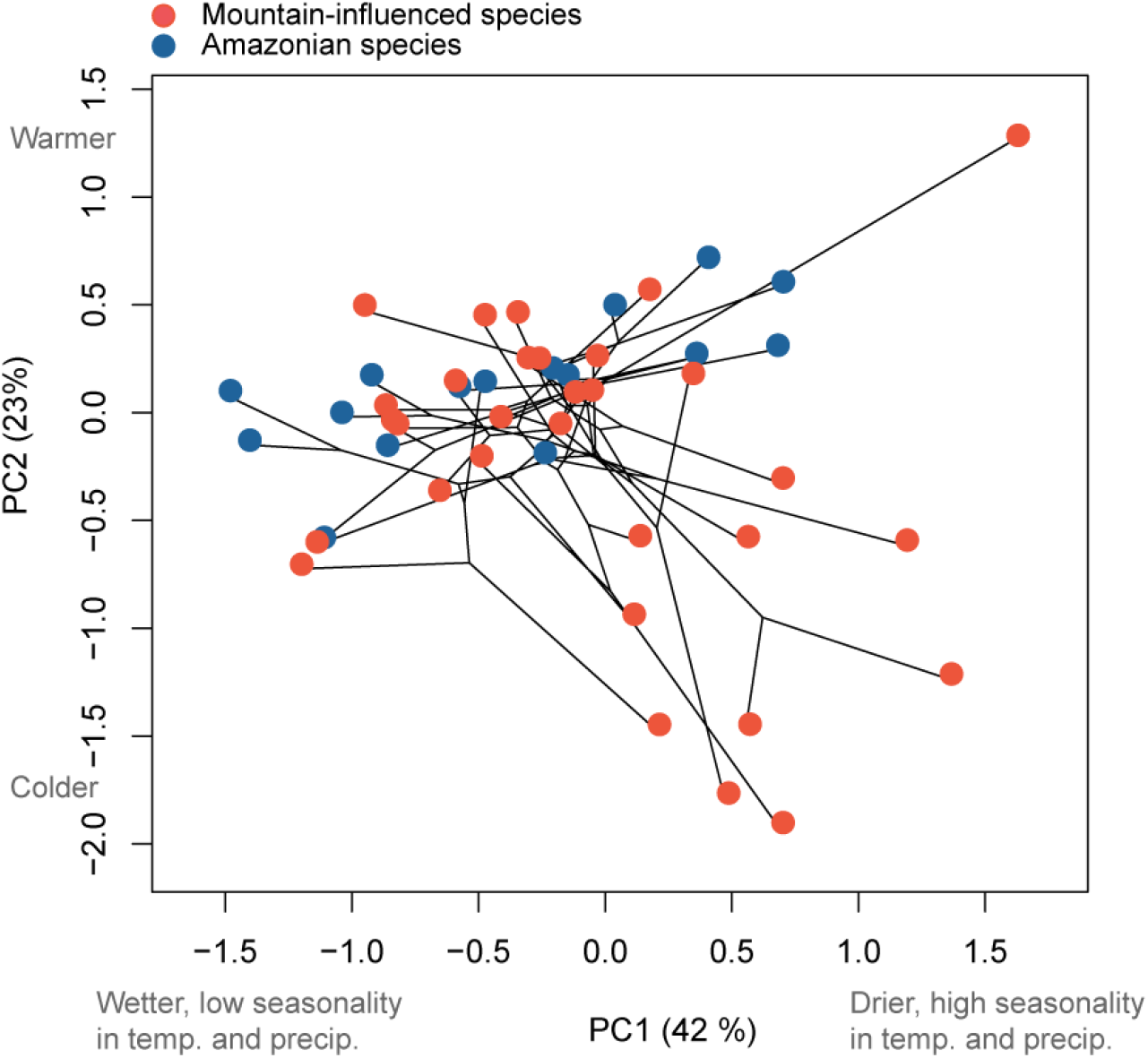
*Costus* phylogeny projected onto the first two principal components for climatic niche. Species are categorized into mountain-influenced (red) and Amazonian (blue). The variance in PC2, but not PC1, is significantly different between categories (see main text).

### EVOLUTION OF POLLINATION SYNDROMES IN MOUNTAINS VERSUS THE AMAZON

The most likely scenario for the evolution of pollination syndromes in our phylogeny involves 11 shifts from orchid bee- to hummingbird-pollination, with seven shifts happening in recent divergence events and four along internal branches of the phylogeny. We find no significant difference in the proportion of hummingbird-pollinated taxa between mountain-influenced and Amazonian taxa (21 out of 39 taxa v. 5 out of 16 taxa; *X*^2^ (1, N = 55) = 1.51, *P* = 0.220; Fig. S6A). We find no evidence of correlated evolution of pollination syndrome and geographic region (Pagel’s test, LR = 0.38, *P* = 0.984). Similarly, we find no significant difference in the frequency of shifts to hummingbird pollination in ancestors characterized as mountain-influenced or Amazonian (8 shifts along 81 branches v. 3 shifts along 28 branches; *X^2^* (NA, N = 109) = 0.02, *P* = 1; Fig. S6B).

### MOUNTAIN-INFLUENCED AND AMAZONIAN SISTER PAIR COMPARISON

Climate niche divergence does not differ between pair types. Thirty-one sister pairs were inferred from the bootstrap replicates of the phylogenetic analysis, and 24 have enough geographic and taxonomic information for comparative analyses (Table S4, Fig. S7). Of these, 15 pairs are identified as mountain-influenced and 9 pairs as Amazonian. We test for niche equivalency for 22 out of our 24 pairs (2 taxa have less than 5 occurrences) rejecting niche equivalency for 60% of sister pairs (Table S4, Fig. S8). We find no difference in the proportion of mountain-influenced and Amazonian pairs with equivalent niches (X^2^ (1, N=22) = 14.29, P = 0.217). Climatic niche divergence is on average 46% greater for mountain influenced pairs relative to Amazonian pairs; however, this difference is not significant (weighted t-test, t = - 1.47^20.2df^, P = 0.157; Figs. 6, 7).

**FIGURE. 6.**
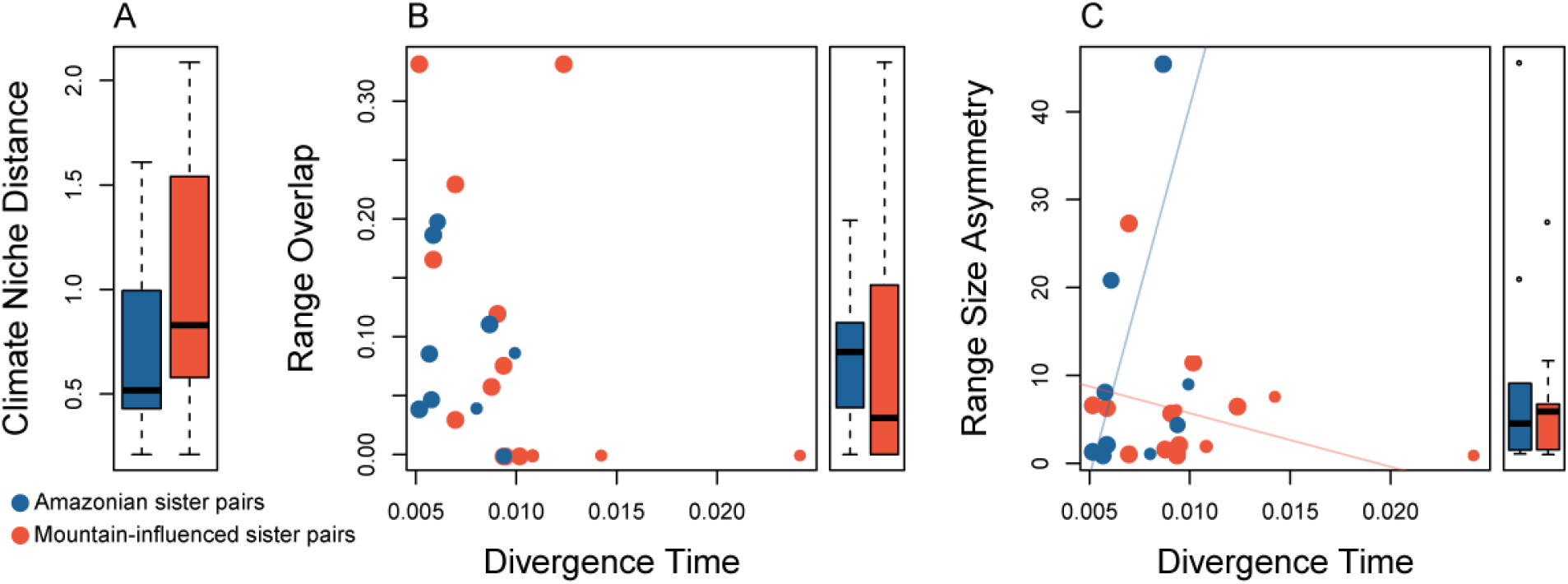
Mountain-influenced and Amazonian sister pair comparisons. A) Climate niche Euclidean distance by region. B) Range overlap v. divergence time (Mya) by region with marginal boxplots indicating differences in range overlap by region. C) Range size asymmetry v. divergence time (Mya) by region, with marginal boxplots indicating differences in range size asymmetry by region. Range overlap and asymmetry were calculated on a grid of 0.05 decimal degrees. In scatterplots, every pair is represented by one dot and its size is proportional to its bootstrap support. Regression lines based on predicted values are included for model effects with uncorrected P<0.05. See main text for statistical results.

**FIGURE. 7.**
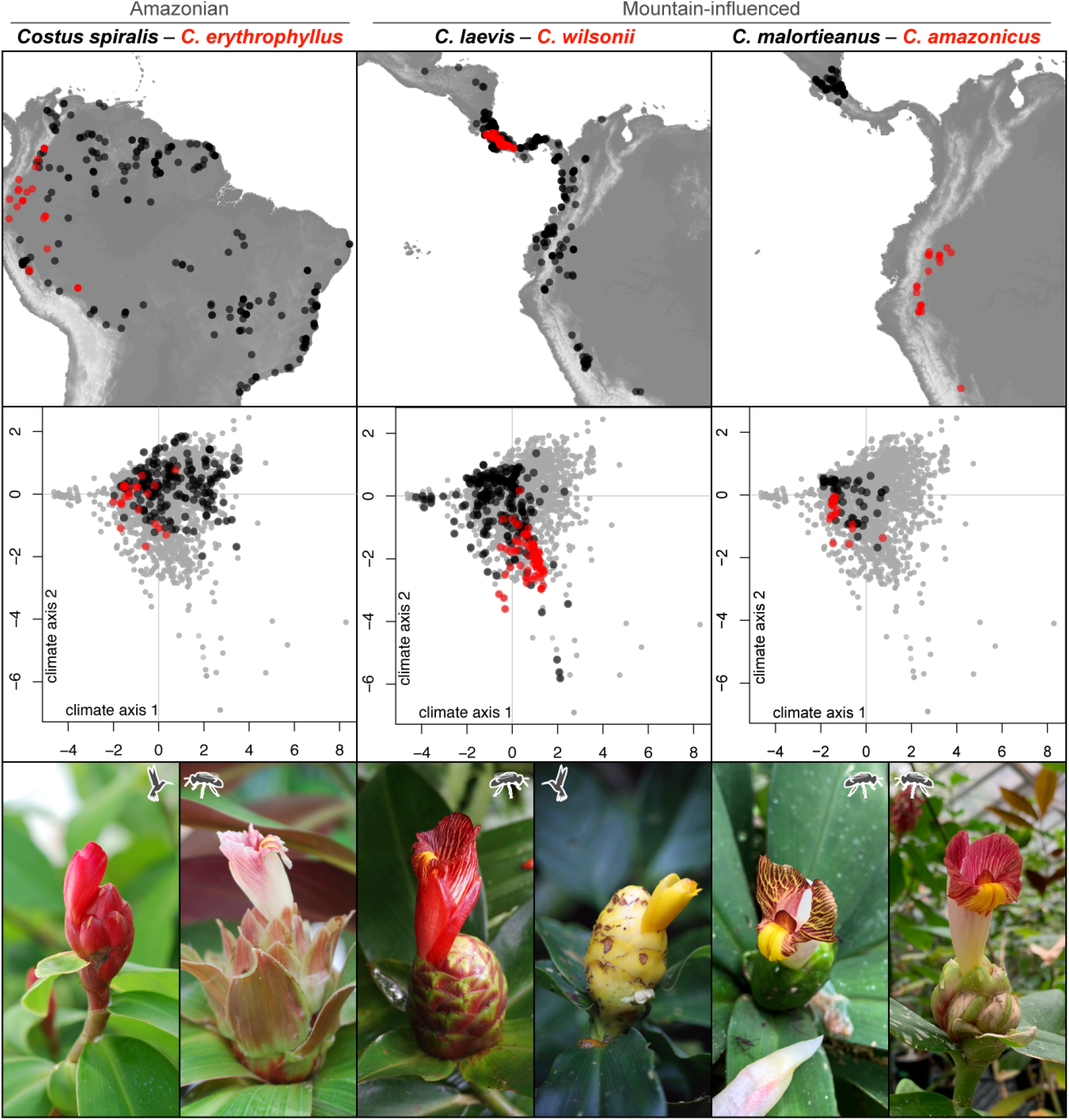
Range maps, climate niches, and inflorescence photos of representative sister species pairs. Left: an Amazonian pair. Center and right: mountain influenced pairs. Top row: occurrences in the map. Central row: comparisons between the climate niches of the sister species (PC axes correspond to Fig. 5). Photo credits from left to right: *Costus spiralis* and *C. erythrophyllus* by KMK, *C. laevis* by R. Maguiña, *C. wilsonii* by P. Juarez, *C. malortieanus* by DLG, and *C. amazonicus* by KMK.

We find no support for the hypothesis that range overlap differs in the mountains versus lowlands. Overall, there is generally little range overlap for sister pairs and no evidence that range overlap varies by region. Average range overlap is 0.09 for mountain influenced and Amazonian pairs at the fine spatial scale (weighted t-test, t = −0.56^22df^, P = 0.584; Figs. 6, 7). Range overlap is not predicted by divergence time, region, or their interaction (Fig. 6; linear model: divergence time F = 0.50^1,20df^, P = 0.487; region F = 0.78^1,20df^, P = 0.386; divergence time by region F = 0.006^1,20df^, P = 0.937). We note that there are two outlier data points with range overlap >0.3 (*C. montanus – C. barbatus, C. ricus – C. scaber*), breaking model assumptions. When those points are removed from the analysis, range overlap decreases with divergence time, consistent with parapatric speciation, but is not predicted by region or their interaction (divergence time F = 6.63^1,18df^, P = 0.019; region F = 0.44^1,18df^, P = 0.515; divergence time by region F = 1.35^1,18df^, P = 0.261). We urge caution interpreting this result, since there is no biological justification for excluding the two outliers.

We find limited support for a budding model of speciation being more common in the mountains than lowlands. Range size asymmetry is greater for younger relative to older mountain-influenced pairs, while the opposite is observed for Amazonian pairs (Fig. 6; GLM: divergence time X^2^ = 0.039, P = 0.843; region X^2^ = 1.747, P = 0.186; divergence time by region X^2^ = 4.34, P = 0.037). In light of multiple comparisons between sister pair types, we treat this result with caution. Note that we present only range overlap and asymmetry results for the fine spatial scale above (~33 km2). Results at the course spatial scale were qualitatively similar for all tests (Fig. S9). Finally, we assess evidence for nested phylogenetic relationships between four taxon pairs, three mountain-influenced and one Amazonian (*C. scaber* in Central America – *C. ricus, C. pulverulentus – C*. sp. nov.18020/18049, *C. laevis – C. wilsonii*, and *C. scaber* in South America – *C. spicatus*, respectively). The first two of those pairs show paraphyly consistent with budding speciation, whereas the latter two are reciprocally monophyletic, although they all show substantial range asymmetry (Fig. 2, Table S4).

## Discussion

Mountains are associated with exceptionally high plant diversity in the Neotropics, with tree species richness peaking in the forests nestled at the eastern base of the Northern Andes (ter Steege et al. 2010) and some of the fastest known plant radiations on earth occur in high elevation Neotropical habitats (Drummond et al. 2012; Madriñán et al. 2013, Uribe-Convers and Tank 2015, Lagomarsino et al. 2016; Vargas et al. 2017, Contreras-Ortiz et al. 2018; Morales-Briones et al. 2018). We see a similar pattern in Neotropical *Costus*, with species richness positively correlated with topographic complexity across its geographic range in Central and South America.

Despite the important contribution of tropical mountains to the latitudinal diversity gradient, the mechanisms underlying this pattern remain unclear. Gentry (1982) hypothesized that recent mountain uplift in the Andes and southern Central America promoted rapid diversification of plants with short generation times (e.g., herbs, shrubs, vines and epiphytes) by providing strong ecogeographic gradients in both climate and pollinators, especially hummingbirds. In contrast, Janzen (1967; reviewed in Sheldon et al. 2018) proposed that the lack of strong temperature seasonality in the tropics leads to narrow physiological tolerances and greater potential for allopatric speciation due to topographic dispersal barriers in mountains. The relative importance of these two mechanisms in driving species richness is unclear—investigation requires understanding species-level relationships in clades that span both montane and lowland environments, and until now we have generally been left comparing Amazonian trees (e.g., Fine et al. 2005; Vargas and Dick 2020) to high elevation shrubs and herbs (e.g., Contreras-Ortiz et al. 2017; Vargas and Simpson 2019). *Costus* provides an opportunity to use a species-level phylogeny to examine possible speciation mechanisms in a clade that spans lowlands to cloud forests, albeit for a single herbaceous life form.

We first examine ecogeographic divergence caused by macroclimatic conditions and adaptation to different functional groups of pollinators: orchid bees and hummingbirds. We find that both mountain-influenced and Amazonian pairs experience climate niche divergence at similar frequencies (Fig. 6. Table S4), with only a marginal trend of greater climatic niche divergence in mountain-influenced pairs, a remarkable result given the steep gradients in climate in tropical mountains. Nevertheless, montane species occupy a significantly greater amount of climatic niche space overall, primarily due to expansion into cooler environments (Fig. 5). Taken together, our results suggest that climatic divergence occurs in both mountain-influenced and Amazonian pairs, and that mountain-influenced taxa occupy the greater temperature variation the mountains offer (Rahbek et al. 2019a). Despite hummingbird pollination being common in mountain-influenced species, it is not proportionally more common than in Amazonian species. Moreover, we observe similar proportions of pollination shifts throughout the tree when ancestors are categorized as mountain-influenced or Amazonian based on our biogeographic reconstruction (Fig. 4,S6). Thus, while hummingbird pollination may be an important driver of diversification in the mountain-influenced pairs, our results show that it is similarly important in lowland Amazonian pairs.

If mountains serve as dispersal barriers and cause long-lasting allopatric separation, we predicted that mountain-influenced pairs would have less range overlap than Amazonian pairs. We find instead that there is generally little range overlap for sister species, regardless of whether they are mountain-influenced or Amazonian (Fig. 6). Although slightly more mountain-influenced than Amazonian pairs have complete allopatric separation (e.g., *C. amazonicus – C. malortieanus*; Fig. 7, right panel), there is no significant difference between regions. This result may simply reflect the importance of geographic isolation for most, if not all, speciation. Our results contrast with a previous study in *Costus*, which used species distribution models to predict co-occurrence across all nodes in the phylogeny and found extensive sympatry (André et al. 2016). Species distribution models may lead to dramatic overestimates of actual co-occurrence (Guisan and Rahbek 2011), particularly in topographically complex landscapes where dispersal is likely limited. Furthermore, because geographic signatures of speciation erode over time as ranges expand, contract, and shift (Fitzpatrick and Turelli 2006), the use of sister species comparisons, rather than all nodes in the phylogeny is more likely to yield information regarding speciation itself.

Additionally, we examined evidence for budding speciation, which may be common when speciation is driven by topographic dispersal barriers (Anacker and Strauss 2014; Grossenbacher et al. 2014). Indirect evidence of budding speciation could come from range size asymmetry decreasing over time-since-divergence, since derivative species should start from small marginal populations. We find a weak pattern of this being the case in mountain-influenced, but not Amazonian pairs (Fig. 6C). Further evidence for budding speciation could come from geographically intensive phylogenetic sampling of sister pairs showing that the smaller-ranged species is nested within the widespread, paraphyletic progenitor (Baldwin 2005). While we find support for this pattern in two mountain-influenced pairs, our phylogenetic sampling was not extensive enough to make statistical comparisons with Amazonian taxa. Thus, while budding speciation likely occurs in *Costus*, our results are not sufficient to draw robust conclusions about the prevalence of budding speciation in the mountains or differences in the frequency of budding speciation between regions. In that sense, our results contrast with the clear patterns of budding speciation found in other plant and animal biodiversity hotspots (Anacker and Strauss 2014; Grossenbacher et al. 2014; Gaboriau et al. 2018).

We see examples of how climatic niche divergence, range overlap and asymmetry, and pollination shifts can occur in both mountain-influenced and Amazonian pairs (Fig. 7). *Costus laevis* is a widespread bee-pollinated lowland species whose range abuts its restricted (but not phylogenetically nested) montane hummingbird-pollinated sister, *C. wilsonii*, in southern Costa Rica. In this case, speciation may be explained by upslope adaptation to a colder drier environment accompanied by a shift to hummingbird pollination in *C. wilsonii*, perhaps the quintessential ecological specialization model of divergence that Gentry (1982) envisioned for Neotropical mountains (Fig. 1E; Fig. 7 center). In contrast, the lowland Amazonian pair *C. erythrophyllus – C. spiralis* shows comparable levels of range overlap, range asymmetry and climate niche divergence, in this case along PC1 (precipitation and seasonality) rather than PC2 (temperature), as well as a shift in pollination syndrome, but without the direct influence of mountains (Fig. 7 left panel). These examples illustrate that multiple ecogeographic factors can promote lineage splitting in both mountains and lowlands.

How can the similarity in patterns of speciation between mountain-influenced and Amazonian pairs be reconciled with the pattern of increased species richness in mountainous regions we see in *Costus?* This pattern could be driven by other factors contributing to higher rates of diversification in mountainous regions, such as less extinction or more immigration, or to differences in the amount of time *Costus* has spent in mountainous v. lowland regions. Our biogeographic reconstructions suggest the latter—*Costus* likely first established ca. 3 Mya in Central America when the Talamanca Cordillera started to uplift (Driese et al. 2007), and the genus diversified in this region for close to 1.5 My before colonizing the already elevated Andes cordillera (Gregory-Wodzicki 2000) and the Amazon lowlands (Fig. 4). Although we were unable to directly compare diversification rates due to our sample size of species and transitions between regions (Maddison and Fitzjohn 2015), we find that Amazonian sister pairs are significantly younger than mountain-influenced pairs (Fig. S9A). This difference is consistent with the Amazon basin as a region of recent and rapid diversification, and counters the alternative hypothesis of higher rates of diversification in mountainous regions. In general, *Costus* has had more time to diversify in Central America and northwestern South America than in the Amazon basin, without the need for invoking different modes or rates of speciation.

To what extent are our results likely to apply to other Neotropical plant lineages? The role of mountains in the diversification of *Costus*, which is restricted to < ca. 2000 m, may be different from higher elevation tropical montane lineages. Studies of Andean paramo plant groups, which occur on mountain tops above treeline, have found substantial allopatry (*Espeletia*: Diazgranados and Barber 2017; *Linochilus*: Vargas and Simpson 2019) and ecological divergence (Campanulaceae: Lagomarsino et al. 2016; *Lupinus*: Nevado et al. 2016, *Espeletia*: Cortés et al. 2018). Alternatively, the relatively young geological age of Neotropical mountains, including the Andes and Central American Cordillera, may spur rapid diversification simply through the opening of new niche space (Weir and Schluter 2008) and without any consistent difference in speciation modes. Much of the plant species richness and endemism in Neotropical mountains comprises herbs and shrubs with short generation times that are able to take advantage of open mountain niche space quickly (Gentry 1982). In contrast, Neotropical trees typically have their center of diversity in the Amazon lowlands (Gentry 1982) and these lineages can date back to the Paleocene (Dick and Pennington 2019). Finally, diversification studies of plants in the Amazon have found a large role for edaphic ecological specialization (*Protium*: Fine et al. 2014, Misiewicz and Fine 2014), other fine-scale habitat divergence (Gesneriaceae: Roalson and Roberts 2016), and biotic interactions, (*Pitcairnia*: Palma-Silva et al. 2011, *Ruellia*: Tripp and Tsai 2017). Both abiotic and biotic conditions vary across lowland forests, and speciation may typically involve ecogeographic isolation even without the influence of mountains.

Taken together, our study suggests that ecogeographic differentiation and geographic isolation are drivers of speciation in mountainous tropical regions, and that they happen similarly in mountains and tropical lowlands. Although mountains provide a larger overall climate niche landscape (Rahbek et al. 2019a), we find no evidence that speciation modes are fundamentally different. However, we caution that these results may not apply to tropical alpine clades that are able to colonize geologically young and spatially disjunct ecosystems above treeline, and these clades may contribute disproportionately to the species richness and endemism of tropical mountains (Hughes and Atchinson 2015). Further studies with a similar framework to ours are needed to determine whether our conclusions can be generalized across tropical organisms in mountainous and lowland regions. Our study demonstrates the potential of combining species-level phylogenomics with spatial and ecological data to test longstanding hypotheses about diversification in tropical mountains.

## Acknowledgments

This research was funded by NSF Dimensions of Biodiversity grant DEB-1737878 to KMK, DLG, Jennifer Funk (Chapman University), Santiago Ramirez (UC Davis), and Carlos Garcia-Robledo (U Conn). We thank the Organization for Tropical Studies, Tortuguero National Park, Estación Biológica Monteverde, and Estación Tropical La Gamba for facilitating field work in Costa Rica. We also thank the Smithsonian Tropical Research Institute for facilitating field work in Panama. We thank Julia Harencár, Rossana Maguiña, Eleinis Ávila-Lovera, and Pedro Juarez for help with field collecting, and Edgardo Ortiz for assistance with the analysis of the transcriptomic data. All field research was conducted with appropriate collection permits in Costa Rica (M-P-SINAC-PNI-ACAT-026-2018, ACC-PI-027-2018, M-PC-SINAC-PNI-ACLAP-020-2018, INV-ACOSA-076-18, M-PC-SINAC-PNI-ACTo-020-18, and CONAGEBIO R-056-2019-OT and R-058-2019-OT), Panama (SE/AP-13-19), and Peru (Resolución Jefatural N° 007-2017-SERNANP-RNAM-JEF and Resolución de Dirección General N° 231-2017-SERFOR/DGGSPFFS both granted by Servicio Nacional Forestal y de Fauna Silvestre del Estado Peruano). Additional plant samples were graciously contributed by John Kress, Douglas Schemske, Dave Skinner, the National Tropical Botanical Garden, and the MICH, MO, UC, and US herbaria. We thank Jim Velzy, Sylvie Childress, and Sarah Ashlock for greenhouse care of the living *Costus* collection, Hannah Thacker and Jaycee Favela for help with lab work, and Russell White and Andrew Fricker for help with the topographic complexity analysis.

## Data Accessibility Statement

Raw sequence data can be found in GenBank (Tables S1-S2). Data, control files, and custom scripts can be accessed in Dryad https://doi.org/10.5061/dryad.p8cz8w9nk, custom code can also be found at https://bitbucket.org/oscarvargash/costus_speciation

