## Supporting information for "Patterns of speciation are similar across mountainous and lowland regions for a Neotropical plant radiation (Costaceae: *Costus*)"

### Supplementary tables:

**Sup. Table 1**. Samples employed for transcriptome sequencing. All samples were sourced from the greenhouses at the University of California, Santa Cruz.

| **Species** | **Plant number** | **Tissue** | **Buffer** | **GenBank BioSample** |
| --- | --- | --- | --- | --- |
| *Costus guanaiensis* Rusby | 122 | flower bud | RLT | SAMN13824540 |
| *Costus osae* Maas & H.Maas | 7 | leaf meristem and flower bud | RLT | SAMN13824541 |
| *Costus pulverulentus* C.Presl | 151 | leaf meristem | RLC | SAMN13824542 |
| *Costus varzearum* Maas | 15 | leaf meristem | RLC | SAMN13824543 |
| *Costus wilsonii* Maas | 201 | leaf meristem | RLT | SAMN13824544 |

**Sup. Table 2**. Samples employed for exon targeted sequencing. HUPCH: Herbario de la Universidad Peruana Cayetano Heredia, JBNB: Jardin Botanique National de Belgique, MICH: University of Michigan Herbarium, MO: Missouri Botanical Garden Herbarium, MSC: Michigan State University Herbarium, NTBG: National Tropical Botanical Garden, TWC: Tom Wood personal collection, UBG: Utrecht Botanic Gardens, UC: Herbarium University of California - Berkeley, UCB Botanical Garden University of California - Berkeley, UCONN Greenhouses University of Connecticut, UCSC: Greenhouses University of California - Santa Cruz, US: United States National Herbarium.

| **Species** | **voucher** | **Coll.** | **Country** | **GB BioSample** | **Code** |
| --- | --- | --- | --- | --- | --- |
| *Costus acreanus* (Loes.) Maas | Skinner R3265 | UC | Peru | SAMN13811711 | C_acre19245 |
| *Costus acreanus* (Loes.) Maas | Rimanchi 10995 | US | Peru | SAMN13811712 | C_acre19253 |
| *Costus afer* Ker Gawl. | 1968GR00199, Wit s.n. (1961) | UBG | Ivory Coast | SAMN13811715 | C_afer98068 |
| *Costus allenii* Maas | Kay 0314 | MSC | Panama | SAMN13811723 | C_alle98050 |
| *Costus amazonicus* (Loes.) J.F.Macbr. | Skinner R3079 | UCSC | Ecuador | SAMN13811724 | C_amaz18009 |
| *Costus amazonicus* (Loes.) J.F.Macbr. | Skinner R3198 | UCSC | Ecuador | SAMN13811725 | C_amaz19020 |
| *Costus amazonicus* subsp. *krukovii* Maas | 1972GR00353, Prance 12826 | UBG | Brazil | SAMN13811726 | C_amaz98072 |
| *Costus arabicus* L. | 1995GRO1263, Maas s.n. | UBG | Brazil | SAMN13811727 | C_arab98080 |
| *Costus asplundii* (Maas) Maas | Maguiña 336 | HUPCH | Peru | SAMN13811728 | C_aspl18058 |
| *Costus barbatus* Suess. | Skinner R3213 | NTBG | Costa Rica | SAMN13811729 | C_barb19242 |
| *Costus bracteatus* Rowlee | Harencar 18-005 (EA0041) | UC | Costa Rica | SAMN13811731 | C_brac18030 |
| *Costus chartaceus* Mass | Kress 90-3124 | UCONN | Colombia | SAMN13811732 | C_char19269 |
| *Costus comosus* var. *bakeri* (K.Schum.) Maas | Skinner R3315 | NTBG | Mexico | SAMN13811736 | C_como19114 |
| *Costus comosus* var. *comosus* (Jacq.) Roscoe | Schemske 032 (T. Wood living coll.) | MSC | unknown | SAMN13811737 | C_como98173 |
| *Costus dirzoi* García-Mend. & G.Ibarra | Ibarra 996 | MO | Mexico | SAMN13811738 | C_dirz19085 |
| *Costus dirzoi* García-Mend. & G.Ibarra | 1980GR00128, Rooden 812 | UBG | Mexico | SAMN13811739 | C_dirz98079 |
| *Costus dubius* (Afzel.) K.Schum. | 1974GR00729, Plowman 4709A | UBG | Peru | SAMN13811742 | C_dubi98074 |
| *Costus dubius* (Afzel.) K.Schum. | Stevenson s.n. (UC-94.1405) | UCB | unknown | SAMN13811740 | C_dubi19268 |
| *Costus dubius* (Afzel.) K.Schum. | US Catalog 1994668 | UCONN | unknown | SAMN13811741 | C_dubi19274 |
| *Costus erythrocoryne* K.Schum. | 1994GR02117, Maas s.n. | UBG | Peru | SAMN13811743 | C_eryt98073 |
| *Costus erythrophyllus* Loes. | Skinner R3306 | NTBG | Colombia | SAMN13811714 | C_AFer19206 |
| *Costus erythrophyllus* Loes. | Kress 94-5379 | US | unknown | SAMN13811734 | C_clav19256 |
| *Costus erythrophyllus* Loes. | Kay 038 | MSC | unknown | SAMN13811744 | C_eryt98168 |
| *Costus erythrophyllus* Loes. | Kay 0339 | MSC | unknown | SAMN13811745 | C_eryt98174 |
| *Costus glaucus* Maas | Skinner R3364 | NTBG | Costa Rica | SAMN13811746 | C_glau19166 |
| *Costus glaucus* Maas | 1974GR00415, Mass 1462 | UBG | Costa Rica | SAMN13811747 | C_glau98077 |
| *Costus guanaiensis* var. *guanaiensis* Rusby | Kay 0317 (UCSC-GH 122) | MSC | Bolivia | SAMN13811752 | C_guan18079 |
| *Costus guanaiensis* var. *guanaiensis* Rusby | Skinner R3140 | NTBG | Panama | SAMN13811753 | C_guan19203 |
| *Costus guanaiensis* var. *guanaiensis* Rusby | Maguiña 326 | HUPCH | Peru | SAMN13811749 | C_guan18056 |
| *Costus guanaiensis* var. *guanaiensis* Rusby | Maguiña 338 | HUPCH | Peru | SAMN13811750 | C_guan18059 |
| *Costus guanaiensis* var*. macrostrobilus* (K.Schum.) Maas | Kay 0319 (UCSC-GH 41) | MSC | Panama | SAMN13811751 | C_guan18077 |
| *Costus guanaiensis* var. *tarmicus* (Loes.) Maas | Skinner s.n. | UCSC | Brazil | SAMN13811748 | C_guan18002 |
| *Costus guanaiensis* var. *tarmicus* (Loes.) Maas | Skinner R3323 | NTBG | Ecuador | SAMN13811717 | C_AFgu19243 |
| *Costus laevis* Ruiz & Pav. | Harencar 18-005 (JH0034) | UC | Costa Rica | SAMN13811730 | C_brac18029 |
| *Costus laevis* Ruiz & Pav. | Kay 0320 | MSC | Costa Rica | SAMN13811754 | C_laev18031 |
| *Costus laevis* Ruiz & Pav. | Harencar 18-010 (JH0071) | UC | Costa Rica | SAMN13811755 | C_laev18032 |
| *Costus laevis* Ruiz & Pav. | Harencar 18-016 (EA0087) | UC | Costa Rica | SAMN13811756 | C_laev19191 |
| *Costus laevis* Ruiz & Pav. | Avila 0068 (EA0070) | UC | Costa Rica | SAMN13811792 | C_ricu19218 |
| *Costus laevis* Ruiz & Pav. | Kay 0310 (UCSC-GH 54) | MSC | Panama | SAMN13811757 | C_laevRD299 |
| *Costus lasius* Loes. | Kay 0321 (UCSC-GH 125) | MSC | Panama | SAMN13811758 | C_lasi18080 |
| *Costus lasius* Loes. | Plowman 11668 | NTBG | Peru | SAMN13811759 | C_lasi19117 |
| *Costus leucanthus* Maas | 86GR00130, Maas 6527 | UBG | Colombia | SAMN13811760 | C_leuc98075 |
| *Costus lima* K.Schum. | 75-0400 | JBNB | Colombia | SAMN13811763 | C_lima98165 |
| *Costus lima* K.Schum. | Kay 023 (DG0028) | MSC | Costa Rica | SAMN13811761 | C_lima18035 |
| *Costus lima* K.Schum. | Kay 023 (DG0025) | MSC | Costa Rica | SAMN13811762 | C_lima19228 |
| *Costus longebracteolatus* Maas | s. n. | TWC | Costa Rica | SAMN13811765 | C_long98176 |
| *Costus longebracteolatus* Maas | Maguiña 344 | HUPCH | Peru | SAMN13811764 | C_long18061 |
| *Costus lucanusianus* J.Braun & K.Schum. | 1968GR00220 | UBG | Cameroon | SAMN13811766 | C_luca99103 |
| *Costus malortieanus* H.Wendl. | Kay 0322 (RM0011) | MSC | Costa Rica | SAMN13811767 | C_malo18036 |
| *Costus malortieanus* H.Wendl. | Kay 0322 (JH0028) | MSC | Costa Rica | SAMN13811768 | C_malo19216 |
| *Costus montanus* Maas | Kay 18-002 (JF0001) | UC | Costa Rica | SAMN13811769 | C_mont18037 |
| *Costus montanus* Maas | Kay 18-002 (KK0075) | UC | Costa Rica | SAMN13811770 | C_mont19217 |
| *Costus nitidus* Maas | Skinner R3088 | UCSC | Costa Rica | SAMN13811771 | C_niti18001 |
| *Costus osae* Maas & H.Maas | Harencar 18-009 (JH0096) | UC | Costa Rica | SAMN13811772 | C_osae18038 |
| *Costus osae* Maas & H.Maas | Harencar 18-009 (JH0081) | UC | Costa Rica | SAMN13811773 | C_osae19005 |
| *Costus pictus* D.Don | Norris 13289 | MICH | Mexico | SAMN13811775 | C_pict19284 |
| *Costus pictus* D.Don | 00-5272 | JBNB | unknown | SAMN13811776 | C_pict98160 |
| *Costus plicatus* Maas | Grossenbacher 0026 | UC | Costa Rica | SAMN13811777 | C_plic18039 |
| *Costus plicatus* Maas | Grossenbacher 0026 (PJ0018) | UC | Costa Rica | SAMN13811778 | C_plic19067 |
| *Costus productus* Gleason ex Maas | Kress 94-3708 | US | unknown | SAMN13811779 | C_prod19263 |
| *Costus productus* Gleason ex Maas | Kay 039 (T. Wood living coll. (labeled C. curvibracteatus)) | MSC | unknown | SAMN13811781 | C_prod98170 |
| *Costus productus* Gleason ex Maas | Skinner s.n. | UCONN | unknown | SAMN13811780 | C_prod19281 |
| *Costus pulverulentus* C.Presl | Skinner R3304 | NTBG | Colombia | SAMN13811789 | C_pulv19119 |
| *Costus pulverulentus* C.Presl | Kay 0326 | MSC | Costa Rica | SAMN13811782 | C_pulv18040 |
| *Costus pulverulentus* C.Presl | Harencar 18-004 (EA0030) | UC | Costa Rica | SAMN13811783 | C_pulv18041 |
| *Costus pulverulentus* C.Presl | Harencar 18-012 (PG0017) | UC | Costa Rica | SAMN13811784 | C_pulv18042 |
| *Costus pulverulentus* C.Presl | Kay 022 (UCSC-GH 133) | MSC | Costa Rica | SAMN13811785 | C_pulv18081 |
| *Costus pulverulentus* C.Presl | Kay 031 (UCSC-GH 144) | MSC | Mexico | SAMN13811786 | C_pulv18082 |
| *Costus pulverulentus* C.Presl | Kay 0327 (UCSC-GH 146) | MSC | Mexico | SAMN13811787 | C_pulv18083 |
| *Costus pulverulentus* C.Presl | Kay 0328 (UCSC-GH 151) | MSC | Panama | SAMN13811788 | C_pulv18084 |
| *Costus pulverulentus* x *scaber* | Skinner R3157 | NTBG | Panama | SAMN13811790 | C_puxs19208 |
| *Costus ricus* Maas & H.Maas | Avila 0068 (JH0075) | UC | Costa Rica | SAMN13811791 | C_ricu18043 |
| *Costus scaber* Ruiz & Pav. | Kay 0329 (UCSC-GH 160) | MSC | Bolivia | SAMN13811797 | C_scab18085 |
| *Costus scaber* Ruiz & Pav. | Kay 032 (UCSC-GH 162) | MSC | Bolivia | SAMN13811798 | C_scab18086 |
| *Costus scaber* Ruiz & Pav. | not collected (la Gamba, Finca la Virgen) | NA | Costa Rica | SAMN13811719 | C_AFsc18044 |
| *Costus scaber* Ruiz & Pav. | Kay 0330 | MSC | Costa Rica | SAMN13811794 | C_scab18045 |
| *Costus scaber* Ruiz & Pav. | Harencar 18-008 (PJ0004) | UC | Costa Rica | SAMN13811795 | C_scab18046 |
| *Costus scaber* Ruiz & Pav. | Kay 021 (UCSC-GH 170) | MSC | Costa Rica | SAMN13811799 | C_scab18087 |
| *Costus scaber* Ruiz & Pav. | Skinner R3215 | NTBG | Guyana | SAMN13811800 | C_scab19167 |
| *Costus scaber Ruiz & Pav.* | Skinner R3220 | NTBG | Guyana | SAMN13811806 | C_spno19122 |
| *Costus scaber* Ruiz & Pav. | Kay 0325 (UCSC-GH 109) | MSC | Panama | SAMN13811796 | C_scab18078 |
| *Costus scaber* Ruiz & Pav. | Maguiña 339 | HUPCH | Peru | SAMN13811720 | C_AFsc18060 |
| *Costus scaber* Ruiz & Pav. | Maguiña 301 | HUPCH | Peru | SAMN13811793 | C_scab18025 |
| *Costus* sp. | Skinner R3308 | NTBG | Colombia | SAMN13811735 | C_como19113 |
| *Costus* sp. | Skinner R3200 | NTBG | Ecuador | SAMN13811716 | C_AFgl19207 |
| *Costus* sp. nov. | Skinner R3226 | NTBG | Colombia | SAMN13811713 | C_AFal19205 |
| *Costus* sp. nov. | Kay 18-001(KK0058) | UC | Costa Rica | SAMN13811721 | C_AFwi18020 |
| *Costus* sp. nov. | Kay 18-001(KK0073) | UC | Costa Rica | SAMN13811722 | C_AFwi18049 |
| *Costus* sp. nov. | Skinner R3314 | NTBG | Mexico | SAMN13811718 | C_AFpi19168 |
| *Costus* sp. nov. | Skinner R3316 | NTBG | Mexico | SAMN13811774 | C_pict19118 |
| *Costus* sp. nov. | Skinner R3130 | UC | Peru | SAMN13811733 | C_clav19248 |
| *Costus spicatus* (Jacq.) Sw. | Skinner R3028 | UC | Dominica | SAMN13811801 | C_spic19251 |
| *Costus spicatus* (Jacq.) Sw. | Kress 02-7143 | UCONN | Dominica | SAMN13811802 | C_spic19272 |
| *Costus spiralis* var. *spiralis* (Jacq.) Roscoe | Kay 033 (UCSC-GH 199) | MSC | Bolivia | SAMN13811803 | C_spir18088 |
| *Costus spiralis* var. *villosus* Maas | Croat 102665 | MO | French Guiana | SAMN13811804 | C_spir19077 |
| *Costus spiralis* var. *villosus* Maas | Skinner R3218 | NTBG | Guyana | SAMN13811805 | C_spir19209 |
| *Costus stenophyllus* Standl. & L.O.Williams | Harencar 18-013 (DG0012) | UC | Costa Rica | SAMN13811807 | C_sten18048 |
| *Costus stenophyllus* Standl. & L.O.Williams | Harencar 18-013 (DG0008) | UC | Costa Rica | SAMN13811808 | C_sten19227 |
| *Costus vargasii* Maas & H.Maas | Skinner R3264 | NTBG | Peru | SAMN13811809 | C_varg19210 |
| *Costus varzearum* Maas | 1971GR00153, Prance 12064 | UBG | Brazil | SAMN13811811 | C_varz98076 |
| *Costus varzearum* Maas | Skinner R1394 | UC | unknown | SAMN13811810 | C_varz19252 |
| *Costus villosissimus* Jacq. | Skinner R3074 | NTBG | Ecuador | SAMN13811813 | C_vill19169 |
| *Costus villosissimus* Jacq. | Kay 0313 (UCSC-GH 214) | MSC | Panama | SAMN13811812 | C_vill18090 |
| *Costus villosissimus* Jacq. | Kay 0313 | MSC | Panama | SAMN13811814 | C_vill98048 |
| *Costus vinosus* Maas | Dressler s.n. | UCONN | Panama | SAMN13811815 | C_vino19273 |
| *Costus vinosus* Maas | TW s.n. | TWC | Panama | SAMN13811816 | C_vino98171 |
| *Costus wilsonii* Maas | Harencar 18-014 (JH0142) | UC | Costa Rica | SAMN13811817 | C_wils18050 |
| *Costus wilsonii* Maas | Harencar 18-015 (PJ0038) | UC | Costa Rica | SAMN13811818 | C_wils18051 |
| *Costus woodsonii* Maas | Harencar 18-007 (JH0051) | UC | Costa Rica | SAMN13811819 | C_wood18052 |
| *Costus woodsonii* Maas | Harencar 18-007 (JH0069) | UC | Costa Rica | SAMN13811821 | C_wood19232 |
| *Costus woodsonii* Maas | Kay 036 (UCSC-GH 208) | MSC | Panama | SAMN13811820 | C_wood18089 |
| *Costus zamoranus* Steyerm. | Skinner R3325 | NTBG | Ecuador | SAMN13811822 | C_zamo19170 |
| *Costus zingiberoides* J.F.Macbr. | 86-0010 | JBNB | Peru | SAMN13811823 | C_zing98162 |

**Sup. Table 3.** Transcriptomic raw reads downloaded from GenBank for data-mining of clock-like genes used to infer the chronogram for Neotropical *Costus*.

| **Species** | **GenBank SRA** |
| --- | --- |
| *Canna indica* L. | SRX2920016 |
| *Canna* sp. | ERX3508843 |
| *Curcuma longa* L. | ERX2099818 |
| *Dichorisandra thyrsiflora* J.C.Mikan | SRX2920013 |
| *Heliconia* sp. | ERX3508845 |
| *Maranta leuconeura* E.Morren | ERX2099816 |
| *Musa basjoo* Siebold & Zucc. ex Iinuma | SRX2920014 |
| *Orchidantha fimbriata* Holttum | SRX2920015 |
| *Strelitzia reginae* Banks | ERX2099817 |
| *Typha latifolia* L. | SRX1639026 |
| *Zingiber officinale* Roscoe | ERX2099820 |

**Sup. Table 4.** Sister species comparisons. Pairs are categorized, in Amazonian (Amaz). or mountain-influenced (mount.).

|  |  |  |  |  | **Grids occupied** | |  |  |  |  | **Range overlap** | |  | **Niche Equivalency** | **Range Asymmetry fine/coarse** | | **Pollinator** | |
| --- | --- | --- | --- | --- | --- | --- | --- | --- | --- | --- | --- | --- | --- | --- | --- | --- | --- | --- |
| **Sister A** | **Sister B** | **BS** | **Cat.** | **Div. time** | **A** | **B** | **A:pc1** | **A:pc2** | **B:pc1** | **B:pc2** | **coarse** | **fine** | **Eucl. Dist.** | **Schoener's**  **D * p<0.05, **p<0.01** | **fine** | **coarse** | **A** | **B** |
| *C zingiberoides* | *C chartaceus* | 727 | Amaz. | 0.0095 | 4 | 19 | -0.938 | 0.154 | -1.056 | -0.020 | 0 | 0.00 | 0.210 | - | 4.54 | 4.288 | hum | hum |
| *C spiralis* | *C arabicus* | 104 | Amaz. | 0.0081 | 182 | 230 | 0.665 | 0.291 | 0.391 | 0.699 | 0.052 | 0.04 | 0.491 | 0.5147* | 1.198 | 1.152 | hum | bee |
| *C guaniensis var. guanaiensis* | *C acreanus* | 1000 | Amaz. | 0.0053 | 38 | 29 | -0.164 | 0.154 | 0.344 | 0.254 | 0.083 | 0.04 | 0.518 | 0.4269 | 1.523 | 1.588 | bee | bee |
| *C spiralis* | *C erythrophyllus* | 896 | Amaz. | 0.0059 | 182 | 22 | 0.665 | 0.291 | -0.876 | -0.172 | 0.11 | 0.05 | 1.609 | 0.0876** | 8.291 | 9.456 | hum | bee |
| *C amazonicus* | *C arabicus* | 66 | Amaz. | 0.01 | 24 | 230 | -0.592 | 0.101 | 0.391 | 0.699 | 0.13 | 0.09 | 1.151 | 0.1172** | 9.099 | 8.551 | bee | bee |
| *C amazonicus* | *C acreanus* | 923 | Amaz. | 0.0058 | 24 | 29 | -0.592 | 0.101 | 0.344 | 0.254 | 0.131 | 0.09 | 0.949 | 0.2095 | 1.076 | 1.033 | bee | bee |
| *C spicatus* | *C scaber* | 956 | Amaz. | 0.0088 | 9 | 430 | 0.687 | 0.586 | -0.224 | 0.186 | 0.112 | 0.11 | 0.995 | 0.1287 | 45.614 | 41.806 | hum | hum |
| *C erythrocoryne* | *C longebracteolatus* | 994 | Amaz. | 0.006 | 18 | 37 | -1.496 | 0.081 | -1.420 | -0.149 | 0.2 | 0.19 | 0.242 | 0.3985 | 2.314 | 2.335 | hum | bee |
| *C arabicus* | *C varzearum* | 828 | Amaz. | 0.0062 | 230 | 11 | 0.391 | 0.699 | 0.022 | 0.479 | 0.222 | 0.20 | 0.430 | 0.0355** | 21.016 | 21.945 | bee | bee |
| *C erythrothyrsus* | *C sp nov* | 1000 | Amaz. | 0.006 | - | - | - | - | - | - | - | - | - | - | - | - | hum | bee |
| *C stenophyllus* | *C vinosus* | 7 | Mount. | 0.0242 | 8 | 7 | -0.491 | 0.433 | -0.068 | 0.084 | 0 | 0.00 | 0.549 | 0.1752 | 1.002 | 1.402 | hum | bee |
| *C osae* | *C comosus var. comosus* | 1000 | Mount. | 0.0095 | 18 | 9 | -0.361 | 0.446 | -0.428 | -0.040 | 0 | 0.00 | 0.490 | 0.1111 | 1.101 | 1.498 | hum | hum |
| *C malortieanus* | *C amazonicus* | 1000 | Mount. | 0.0096 | 61 | 18 | -0.856 | -0.054 | -1.126 | -0.600 | 0 | 0.00 | 0.609 | 0.2866* | 2.298 | 2.086 | bee | bee |
| *C woodsonii* | *C scaber* | 44 | Mount. | 0.0143 | 56 | 430 | -0.966 | 0.478 | -0.224 | 0.186 | 0 | 0.00 | 0.797 | 0.1461** | 7.68 | 8.925 | hum | hum |
| *C guanaiensis var. tarmicus* | *C guanaiensis var. macrostrobilus* | 250 | Mount. | 0.0109 | 51 | 89 | 0.548 | -0.596 | -0.047 | 0.244 | 0 | 0.00 | 1.029 | 0.2558** | 2.066 | 2.24 | bee | bee |
| *C lima* | *C comosus var. bakeri* | 987 | Mount. | 0.0103 | 104 | 12 | -0.321 | 0.234 | 0.685 | -0.324 | 0 | 0.00 | 1.151 | 0.188 | 11.668 | 12.267 | hum | hum |
| *C zamoranus* | *C longebracteolatus* | 6 | Mount. | 0.0094 | 7 | 37 | 0.197 | -1.468 | -1.420 | -0.149 | 0 | 0.00 | 2.087 | 0.0364* | 6.177 | 8.764 | bee | bee |
| *C allenii* | *C bracteatus* | 1000 | Mount. | 0.0071 | 49 | 57 | -0.193 | -0.072 | -0.883 | 0.014 | 0.08 | 0.03 | 0.695 | 0.199* | 1.245 | 1.394 | bee | bee |
| *C productus* | *C vargasii* | 1000 | Mount. | 0.0089 | 32 | 18 | -0.253 | -0.207 | 1.175 | -0.612 | 0.059 | 0.06 | 1.485 | 0.077* | 1.788 | 1.669 | hum | hum |
| *C plicatus* | *C nitidus* | 1000 | Mount. | 0.0095 | 23 | 13 | -0.607 | 0.128 | -0.670 | -0.381 | 0.091 | 0.08 | 0.513 | 0.2274 | 1.31 | 1.183 | hum | hum |
| *C laevis* | *C wilsonii* | 1000 | Mount. | 0.0092 | 237 | 68 | -0.504 | -0.222 | 0.685 | -1.923 | 0.115 | 0.12 | 2.076 | 0.1094** | 5.889 | 6.589 | bee | hum |
| *C asplundii* | *C zamoranus* | 994 | Mount. | 0.006 | 40 | 7 | -1.215 | -0.724 | 0.197 | -1.468 | 0.25 | 0.17 | 1.596 | 0.1224 | 6.508 | 8.76 | bee | bee |
| *C sp nov* | *C pulverulentus* | 1000 | Mount. | 0.0071 | 17 | 421 | 0.471 | -1.786 | 0.331 | 0.161 | 0.2 | 0.23 | 1.951 | 0.0496** | 27.5 | 31.264 | hum | hum |
| *C ricus* | *C scaber* | 1000 | Mount. | 0.0053 | 27 | 186 | -0.276 | 0.232 | -0.134 | 0.075 | 0.6 | 0.33 | 0.211 | 0.1138** | 6.815 | 8.343 | hum | hum |
| *C montanus* | *C barbatus* | 1000 | Mount. | 0.0125 | 31 | 4 | 0.555 | -1.467 | 1.350 | -1.233 | 0.333 | 0.33 | 0.829 | - | 6.663 | 6.664 | hum | hum |
| *C chartaceus* | *C sp nov* | 273 | Mount. | 0.0139 | - | - | - | - | - | - | - | - | - | - | - | - | bee | hum |
| *C glaucus* | *C sp nov* | 1000 | Mount. | 0.0101 | - | - | - | - | - | - | - | - | - | - | - | - | bee | bee |
| *C guanaiensis* | *C amazonicus* | 1000 | Mount. | 0.005 | - | - | - | - | - | - | - | - | - | - | - | - | bee | bee |
| *C guanaiensis var. tarmicus* | *Costus sp.* | 750 | Mount. | 0.0068 | - | - | - | - | - | - | - | - | - | - | - | - | bee | bee |
| *C pictus* | *C dirzoi* | 1000 | Mount. | 0.0098 | - | - | - | - | - | - | - | - | - | - | - | - | bee | bee |
| *C sp* | *C villosissimus* | 1000 | Mount. | 0.0054 | - | - | - | - | - | - | - | - | - | - | - | - | hum | bee |
| *C lucanusianus* | *C afer* | 1000 | Outgroup | - | - | - | - | - | - | - | - | - | - | - | - | - | - | - |

**Sup. Table 5.** Sample statistics: gene-regions, number of characters in supermatrix, proportion of gene-regions, and percentage of columns with data in the supermatrix.

| **Sample** | **# Regions** | **# characters** | **% regions** | **% occupancy** |
| --- | --- | --- | --- | --- |
| Costus_acreanus_19245 | 727 | 1,398,370 | 96 | 95 |
| Costus_acreanus_19253 | 510 | 863,525 | 67 | 59 |
| Costus_afer_98068 | 698 | 1,335,436 | 92 | 91 |
| Costus_allenii_98050 | 745 | 1,446,971 | 99 | 98 |
| Costus_amazonicus_19020 | 740 | 1,433,735 | 98 | 97 |
| Costus_amazonicus_GUAL_18009 | 740 | 1,439,403 | 98 | 98 |
| Costus_amazonicus_subsp_krukovii_98072 | 723 | 1,386,772 | 96 | 94 |
| Costus_arabicus_98080 | 746 | 1,448,978 | 99 | 98 |
| Costus_asplundii_18058 | 749 | 1,451,016 | 99 | 98 |
| Costus_barbatus_19242 | 743 | 1,442,155 | 98 | 98 |
| Costus_bracteatus_18030 | 735 | 1,419,812 | 97 | 96 |
| Costus_chartaceus_19269 | 747 | 1,454,236 | 99 | 99 |
| Costus_comosus_var_bakeri_19114 | 735 | 1,413,841 | 97 | 96 |
| Costus_comosus_var_comosus_98173 | 732 | 1,414,961 | 97 | 96 |
| Costus_dirzoi_19085 | 377 | 579,039 | 50 | 39 |
| Costus_dirzoi_98079 | 705 | 1,348,036 | 93 | 91 |
| Costus_dubius_19268 | 696 | 1,311,525 | 92 | 89 |
| Costus_dubius_19274 | 710 | 1,357,835 | 94 | 92 |
| Costus_dubius_98074 | 703 | 1,336,210 | 93 | 91 |
| Costus_erythrocoryne_98073 | 736 | 1,421,756 | 97 | 96 |
| Costus_erythrophyllus_19206 | 743 | 1,442,995 | 98 | 98 |
| Costus_erythrophyllus_19256 | 736 | 1,423,091 | 97 | 96 |
| Costus_erythrophyllus_98168 | 746 | 1,449,080 | 99 | 98 |
| Costus_erythrothyrsus_98174 | 733 | 1,417,635 | 97 | 96 |
| Costus_glaucus_19166 | 738 | 1,426,474 | 98 | 97 |
| Costus_glaucus_98077 | 745 | 1,450,098 | 99 | 98 |
| Costus_guanaiensis_var_guanaiensis_CA_19203 | 744 | 1,443,658 | 98 | 98 |
| Costus_guanaiensis_var_guanaiensis_SA_18056 | 743 | 1,436,568 | 98 | 97 |
| Costus_guanaiensis_var_guanaiensis_SA_18059 | 744 | 1,444,620 | 98 | 98 |
| Costus_guanaiensis_var_guanaiensis_SA_18079 | 748 | 1,456,716 | 99 | 99 |
| Costus_guanaiensis_var_macrostrobilus_18077 | 744 | 1,441,932 | 98 | 98 |
| Costus_guanaiensis_var_tarmicus_18002 | 743 | 1,440,990 | 98 | 98 |
| Costus_guanaiensis_var_tarmicus_19243 | 744 | 1,440,518 | 98 | 98 |
| Costus_laevis_18029 | 737 | 1,429,846 | 97 | 97 |
| Costus_laevis_18031 | 735 | 1,417,123 | 97 | 96 |
| Costus_laevis_18032 | 744 | 1,446,820 | 98 | 98 |
| Costus_laevis_19191 | 744 | 1,436,958 | 98 | 97 |
| Costus_laevis_19218 | 741 | 1,432,552 | 98 | 97 |
| Costus_laevis_RD299 | 740 | 1,430,345 | 98 | 97 |
| Costus_lasius_18080 | 745 | 1,447,947 | 99 | 98 |
| Costus_lasius_19117 | 732 | 1,413,879 | 97 | 96 |
| Costus_leucanthus_98075 | 705 | 1,357,648 | 93 | 92 |
| Costus_lima_18035 | 733 | 1,420,662 | 97 | 96 |
| Costus_lima_19228 | 746 | 1,450,500 | 99 | 98 |
| Costus_lima_98165 | 744 | 1,448,795 | 98 | 98 |
| Costus_longebracteolatus_18061 | 736 | 1,418,238 | 97 | 96 |
| Costus_longebracteolatus_98176 | 733 | 1,410,870 | 97 | 96 |
| Costus_lucanusianus_99103 | 740 | 1,432,373 | 98 | 97 |
| Costus_malortieanus_18036 | 734 | 1,418,207 | 97 | 96 |
| Costus_malortieanus_19216 | 731 | 1,407,075 | 97 | 95 |
| Costus_montanus_18037 | 738 | 1,423,987 | 98 | 97 |
| Costus_montanus_19217 | 726 | 1,392,694 | 96 | 94 |
| Costus_nitidus_18001 | 741 | 1,431,013 | 98 | 97 |
| Costus_osae_18038 | 735 | 1,417,842 | 97 | 96 |
| Costus_osae_19005 | 736 | 1,419,995 | 97 | 96 |
| Costus_pictus_19284 | 527 | 905,843 | 70 | 61 |
| Costus_pictus_98160 | 700 | 1,320,456 | 93 | 90 |
| Costus_plicatus_18039 | 745 | 1,_450_,976 | 99 | 98 |
| Costus_plicatus_19067 | 744 | 1,446,258 | 98 | 98 |
| Costus_productus_19263 | 712 | 1,362,616 | 94 | 92 |
| Costus_productus_19281 | 737 | 1,419,082 | 97 | 96 |
| Costus_productus_98170 | 743 | 1,444,318 | 98 | 98 |
| Costus_pulverulentus_18040 | 733 | 1,414,809 | 97 | 96 |
| Costus_pulverulentus_18041 | 739 | 1,432,867 | 98 | 97 |
| Costus_pulverulentus_18042 | 745 | 1,446,150 | 99 | 98 |
| Costus_pulverulentus_18081 | 740 | 1,441,154 | 98 | 98 |
| Costus_pulverulentus_18082 | 738 | 1,431,859 | 98 | 97 |
| Costus_pulverulentus_18083 | 745 | 1,440,431 | 99 | 98 |
| Costus_pulverulentus_18084 | 749 | 1,460,259 | 99 | 99 |
| Costus_pulverulentus_19119 | 745 | 1,450,914 | 99 | 98 |
| Costus_pulverulentus_x_scaber_19208 | 745 | 1,450,383 | 99 | 98 |
| Costus_ricus_18043 | 744 | 1,448,428 | 98 | 98 |
| Costus_scaber_18025 | 729 | 1,391,398 | 96 | 94 |
| Costus_scaber_18044 | 742 | 1,443,647 | 98 | 98 |
| Costus_scaber_18045 | 746 | 1,449,694 | 99 | 98 |
| Costus_scaber_18046 | 745 | 1,448,356 | 99 | 98 |
| Costus_scaber_18060 | 739 | 1,437,076 | 98 | 97 |
| Costus_scaber_18078 | 741 | 1,438,794 | 98 | 98 |
| Costus_scaber_18085 | 749 | 1,455,265 | 99 | 99 |
| Costus_scaber_18086 | 735 | 1,414,574 | 97 | 96 |
| Costus_scaber_18087 | 745 | 1,451,531 | 99 | 98 |
| Costus_scaber_19167 | 746 | 1,452,681 | 99 | 98 |
| Costus_sp_19113 | 738 | 1,433,483 | 98 | 97 |
| Costus_scaber_19122 | 740 | 1,427,399 | 98 | 97 |
| Costus_sp_19207 | 746 | 1,446,134 | 99 | 98 |
| Costus_sp_nov_18020 | 745 | 1,446,557 | 99 | 98 |
| Costus_sp_nov_18049 | 742 | 1,438,733 | 98 | 98 |
| Costus_sp_nov_19118 | 740 | 1,431,934 | 98 | 97 |
| Costus_sp_nov_19168 | 745 | 1,445,079 | 99 | 98 |
| Costus_sp_nov_19205 | 741 | 1,438,462 | 98 | 98 |
| Costus_sp_nov_19248 | 727 | 1,409,731 | 96 | 96 |
| Costus_spicatus_19251 | 741 | 1,439,651 | 98 | 98 |
| Costus_spicatus_19272 | 741 | 1,439,219 | 98 | 98 |
| Costus_spiralis_var_spiralis_18088 | 741 | 1,431,687 | 98 | 97 |
| Costus_spiralis_var_villosus_19077 | 550 | 970,858 | 73 | 66 |
| Costus_spiralis_var_villosus_19209 | 736 | 1,425,424 | 97 | 97 |
| Costus_stenophyllus_18048 | 745 | 1,450,876 | 99 | 98 |
| Costus_stenophyllus_19227 | 741 | 1,439,853 | 98 | 98 |
| Costus_vargasii_19210 | 747 | 1,442,985 | 99 | 98 |
| Costus_varzearum_19252 | 735 | 1,420,283 | 97 | 96 |
| Costus_varzearum_98076 | 711 | 1,379,738 | 94 | 94 |
| Costus_villosissimus_18090 | 750 | 1,459,503 | 99 | 99 |
| Costus_villosissimus_19169 | 747 | 1,452,747 | 99 | 99 |
| Costus_villosissimus_98048 | 747 | 1,451,911 | 99 | 98 |
| Costus_vinosus_19273 | 748 | 1,454,814 | 99 | 99 |
| Costus_vinosus_98171 | 740 | 1,434,295 | 98 | 97 |
| Costus_wilsonii_18050 | 742 | 1,437,657 | 98 | 97 |
| Costus_wilsonii_18051 | 737 | 1,422,227 | 97 | 96 |
| Costus_woodsonii_18052 | 746 | 1,447,250 | 99 | 98 |
| Costus_woodsonii_18089 | 746 | 1,453,682 | 99 | 99 |
| Costus_woodsonii_19232 | 746 | 1,451,768 | 99 | 98 |
| Costus_zamoranus_19170 | 744 | 1,446,613 | 98 | 98 |
| Costus_zingiberoides_98162 | 667 | 1,236,898 | 88 | 84 |

**Sup. Table 6.** Gene matrices statistics, alignment length and number of samples.

| **Gene** | **Length** | **# taxa** |  | **Gene** | **Length** | **# taxa** |
| --- | --- | --- | --- | --- | --- | --- |
| cluster10001 | 1007 | 112 |  | cluster3055 | 1949 | 111 |
| cluster10040 | 1899 | 104 |  | cluster3060 | 1996 | 113 |
| cluster10069 | 1180 | 113 |  | cluster3086 | 2954 | 111 |
| cluster10091 | 1193 | 112 |  | cluster3091 | 2773 | 111 |
| cluster10102 | 1118 | 111 |  | cluster3102 | 2123 | 112 |
| cluster10126 | 1191 | 113 |  | cluster3114 | 1753 | 112 |
| cluster10184 | 2369 | 106 |  | cluster3134 | 3122 | 110 |
| cluster10227 | 1049 | 113 |  | cluster3219 | 1997 | 113 |
| cluster10245 | 1110 | 113 |  | cluster3235 | 3073 | 110 |
| cluster10322 | 2613 | 107 |  | cluster3281 | 2340 | 109 |
| cluster10331 | 3263 | 109 |  | cluster3286 | 2019 | 112 |
| cluster10345 | 1118 | 113 |  | cluster3311 | 2660 | 110 |
| cluster10385 | 1447 | 113 |  | cluster3327 | 2387 | 107 |
| cluster10412 | 1735 | 104 |  | cluster3376 | 2053 | 113 |
| cluster10421 | 3195 | 105 |  | cluster3377 | 2595 | 112 |
| cluster10439 | 1084 | 113 |  | cluster3387 | 1521 | 112 |
| cluster10492 | 1936 | 105 |  | cluster3389 | 1885 | 111 |
| cluster10518A | 1322 | 112 |  | cluster3407 | 1926 | 113 |
| cluster10522 | 1542 | 112 |  | cluster3414 | 2633 | 113 |
| cluster10554 | 1043 | 113 |  | cluster3416 | 2904 | 113 |
| cluster10609 | 636 | 113 |  | cluster3504 | 1944 | 113 |
| cluster10628 | 1201 | 112 |  | cluster3539 | 3752 | 108 |
| cluster10642 | 1931 | 110 |  | cluster3558 | 1973 | 113 |
| cluster10643 | 2040 | 105 |  | cluster3571 | 3221 | 111 |
| cluster10719 | 1927 | 108 |  | cluster3613 | 1724 | 113 |
| cluster10752 | 760 | 113 |  | cluster3616 | 3038 | 107 |
| cluster10769 | 1612 | 112 |  | cluster3627 | 2730 | 112 |
| cluster10793 | 1760 | 110 |  | cluster3642 | 1777 | 113 |
| cluster10800 | 1890 | 109 |  | cluster3644 | 3734 | 106 |
| cluster10802 | 1023 | 113 |  | cluster3647 | 1946 | 111 |
| cluster10808 | 2478 | 109 |  | cluster3675 | 2800 | 109 |
| cluster10838 | 1949 | 107 |  | cluster3680 | 1680 | 113 |
| cluster10848 | 1191 | 113 |  | cluster3744 | 1991 | 113 |
| cluster10850 | 1866 | 112 |  | cluster3745 | 1889 | 113 |
| cluster10883 | 1666 | 108 |  | cluster3768 | 2994 | 108 |
| cluster10917 | 1284 | 113 |  | cluster3796 | 1859 | 113 |
| cluster10920 | 1859 | 113 |  | cluster3811 | 1634 | 113 |
| cluster10941 | 2076 | 107 |  | cluster3846 | 3995 | 106 |
| cluster10960 | 2085 | 103 |  | cluster3848 | 2782 | 110 |
| cluster1098 | 2054 | 110 |  | cluster3876 | 1939 | 113 |
| cluster11002 | 1821 | 101 |  | cluster3887 | 3084 | 109 |
| cluster11055 | 1874 | 102 |  | cluster3913 | 2505 | 113 |
| cluster11099 | 1230 | 113 |  | cluster3917 | 2785 | 112 |
| cluster11114 | 1536 | 108 |  | cluster3921 | 4153 | 109 |
| cluster11190 | 1942 | 98 |  | cluster3931 | 1693 | 113 |
| cluster11265 | 846 | 113 |  | cluster3939 | 1391 | 113 |
| cluster11269 | 1074 | 112 |  | cluster3963 | 1586 | 113 |
| cluster11274 | 558 | 113 |  | cluster3983 | 2589 | 112 |
| cluster11286 | 920 | 113 |  | cluster3991 | 1983 | 113 |
| cluster11297 | 1744 | 105 |  | cluster3993 | 1923 | 112 |
| cluster11308 | 1972 | 107 |  | cluster4014 | 3231 | 112 |
| cluster11341 | 1585 | 105 |  | cluster4018 | 4345 | 108 |
| cluster11365 | 1071 | 113 |  | cluster4028 | 1424 | 113 |
| cluster11381 | 1962 | 105 |  | cluster4053 | 2974 | 111 |
| cluster11386 | 1775 | 110 |  | cluster4062 | 1939 | 113 |
| cluster11387 | 1892 | 106 |  | cluster4114 | 3068 | 108 |
| cluster11408 | 3249 | 99 |  | cluster4118 | 5442 | 105 |
| cluster11454 | 2766 | 106 |  | cluster4120 | 1608 | 113 |
| cluster11460 | 977 | 113 |  | cluster4123 | 1575 | 112 |
| cluster11483 | 1174 | 111 |  | cluster4129 | 1899 | 112 |
| cluster11547 | 940 | 113 |  | cluster4135 | 1883 | 113 |
| cluster11555 | 1764 | 109 |  | cluster4146 | 1715 | 112 |
| cluster11558 | 2730 | 107 |  | cluster4148 | 2107 | 111 |
| cluster11566 | 1835 | 110 |  | cluster4195 | 1290 | 113 |
| cluster11601 | 2067 | 105 |  | cluster4255 | 2718 | 110 |
| cluster11633 | 1020 | 112 |  | cluster4262 | 1097 | 113 |
| cluster11637 | 287 | 113 |  | cluster4274 | 1853 | 113 |
| cluster11670 | 997 | 113 |  | cluster4298 | 1866 | 113 |
| cluster11675 | 1789 | 109 |  | cluster4325 | 1627 | 113 |
| cluster11684 | 2161 | 110 |  | cluster4331 | 2482 | 112 |
| cluster11709 | 4744 | 81 |  | cluster4336 | 1748 | 112 |
| cluster11712 | 1082 | 113 |  | cluster4338 | 1619 | 113 |
| cluster11715 | 1876 | 110 |  | cluster4349 | 1754 | 111 |
| cluster11737 | 1116 | 112 |  | cluster4381 | 3481 | 103 |
| cluster11740 | 1901 | 106 |  | cluster4382 | 2623 | 112 |
| cluster11757 | 1268 | 112 |  | cluster4385 | 4654 | 106 |
| cluster11787 | 1723 | 111 |  | cluster4407 | 1643 | 112 |
| cluster11809 | 1528 | 108 |  | cluster4409 | 2892 | 108 |
| cluster11815 | 1045 | 112 |  | cluster4416 | 1819 | 113 |
| cluster11819 | 1987 | 107 |  | cluster4424 | 730 | 113 |
| cluster11854 | 2102 | 108 |  | cluster4453 | 1927 | 110 |
| cluster11893 | 1819 | 112 |  | cluster4458 | 1638 | 113 |
| cluster11933 | 833 | 112 |  | cluster4462 | 2542 | 110 |
| cluster11941 | 1967 | 108 |  | cluster4464 | 2728 | 108 |
| cluster11955 | 1426 | 107 |  | cluster4466 | 1760 | 113 |
| cluster11956 | 1986 | 107 |  | cluster4477 | 1480 | 113 |
| cluster11970 | 1056 | 112 |  | cluster4494 | 2894 | 107 |
| cluster1199 | 3349 | 110 |  | cluster4538 | 1902 | 111 |
| cluster11995 | 3140 | 103 |  | cluster4565 | 3041 | 111 |
| cluster12035 | 1180 | 111 |  | cluster4570 | 3244 | 111 |
| cluster12072 | 1120 | 112 |  | cluster4579 | 1802 | 112 |
| cluster12084 | 2651 | 73 |  | cluster4599 | 1647 | 112 |
| cluster12114 | 2592 | 108 |  | cluster4616 | 1828 | 113 |
| cluster12140 | 1027 | 112 |  | cluster4617 | 4492 | 103 |
| cluster12144 | 1458 | 107 |  | cluster4654 | 3759 | 106 |
| cluster12148 | 832 | 113 |  | cluster4664 | 1644 | 113 |
| cluster12154 | 2084 | 107 |  | cluster4667 | 1791 | 113 |
| cluster12160 | 2225 | 105 |  | cluster4675 | 1826 | 113 |
| cluster12228 | 724 | 110 |  | cluster4691 | 2658 | 112 |
| cluster12230 | 3256 | 106 |  | cluster4703 | 2576 | 111 |
| cluster12232 | 3291 | 104 |  | cluster4709 | 1827 | 112 |
| cluster12248 | 2286 | 107 |  | cluster4713 | 1769 | 111 |
| cluster12261 | 1176 | 113 |  | cluster4716 | 2401 | 111 |
| cluster12291 | 1771 | 107 |  | cluster4728 | 1773 | 113 |
| cluster12303 | 1075 | 111 |  | cluster4732 | 1855 | 113 |
| cluster12331 | 1333 | 71 |  | cluster4745 | 2238 | 111 |
| cluster12338 | 1647 | 107 |  | cluster4775 | 1233 | 113 |
| cluster12359 | 2321 | 107 |  | cluster4905 | 1760 | 113 |
| cluster12404 | 948 | 113 |  | cluster4939 | 1707 | 113 |
| cluster12406 | 1003 | 113 |  | cluster4963 | 3502 | 107 |
| cluster12420 | 1717 | 108 |  | cluster4964 | 1604 | 112 |
| cluster12435 | 967 | 113 |  | cluster4975 | 1698 | 112 |
| cluster12455 | 1194 | 112 |  | cluster4985 | 2647 | 111 |
| cluster12457 | 966 | 110 |  | cluster4991 | 1638 | 113 |
| cluster12472 | 1842 | 108 |  | cluster5000 | 1886 | 113 |
| cluster12490 | 609 | 112 |  | cluster5003 | 1135 | 112 |
| cluster12530 | 1079 | 110 |  | cluster5027 | 2179 | 112 |
| cluster12539 | 1805 | 109 |  | cluster5058 | 2209 | 111 |
| cluster12555 | 1550 | 109 |  | cluster5075 | 3772 | 108 |
| cluster12573 | 748 | 110 |  | cluster5080 | 2241 | 112 |
| cluster12594 | 1086 | 113 |  | cluster5131 | 2369 | 112 |
| cluster12609 | 2172 | 107 |  | cluster5136 | 2717 | 108 |
| cluster12659 | 2468 | 108 |  | cluster5144 | 1414 | 113 |
| cluster12669 | 1546 | 112 |  | cluster5160 | 1803 | 113 |
| cluster12673 | 833 | 113 |  | cluster5163 | 1522 | 112 |
| cluster12682 | 1961 | 105 |  | cluster5203 | 2693 | 107 |
| cluster12704 | 1738 | 108 |  | cluster5224 | 1722 | 113 |
| cluster12781 | 1016 | 111 |  | cluster5232 | 1629 | 111 |
| cluster12813 | 1586 | 106 |  | cluster5236 | 1605 | 112 |
| cluster12838 | 1635 | 111 |  | cluster5238 | 3133 | 108 |
| cluster12853 | 959 | 113 |  | cluster5246 | 2183 | 110 |
| cluster12914 | 933 | 113 |  | cluster5253 | 1754 | 112 |
| cluster12941 | 2267 | 102 |  | cluster5282 | 1566 | 111 |
| cluster12992 | 462 | 113 |  | cluster5291 | 1536 | 110 |
| cluster13025 | 1579 | 110 |  | cluster5326 | 4779 | 105 |
| cluster13042 | 941 | 113 |  | cluster5362 | 2770 | 108 |
| cluster13051 | 1686 | 110 |  | cluster5392 | 2984 | 109 |
| cluster13072 | 576 | 113 |  | cluster5407 | 2815 | 106 |
| cluster13077 | 1416 | 111 |  | cluster5440 | 1496 | 113 |
| cluster13084 | 1872 | 106 |  | cluster5452 | 1606 | 113 |
| cluster13128 | 673 | 113 |  | cluster5453 | 2955 | 111 |
| cluster13147 | 2304 | 110 |  | cluster5460 | 2358 | 109 |
| cluster13159 | 926 | 113 |  | cluster5492 | 2869 | 106 |
| cluster13160 | 1239 | 111 |  | cluster5511 | 1309 | 113 |
| cluster13182 | 1064 | 112 |  | cluster5525 | 2362 | 112 |
| cluster13206 | 1132 | 112 |  | cluster5537 | 2368 | 110 |
| cluster13234 | 3373 | 108 |  | cluster5636 | 2855 | 111 |
| cluster13244 | 1696 | 112 |  | cluster5657 | 3887 | 110 |
| cluster13261 | 799 | 111 |  | cluster5660 | 4212 | 109 |
| cluster13352 | 1825 | 110 |  | cluster5670 | 1650 | 111 |
| cluster13419 | 3669 | 95 |  | cluster5683 | 1569 | 113 |
| cluster13440 | 890 | 113 |  | cluster5705 | 2699 | 110 |
| cluster13451 | 1534 | 110 |  | cluster5711 | 1595 | 111 |
| cluster13481 | 459 | 113 |  | cluster5721 | 1737 | 108 |
| cluster13497 | 937 | 113 |  | cluster5725 | 1988 | 110 |
| cluster13554 | 970 | 112 |  | cluster5769 | 1255 | 113 |
| cluster13559 | 2149 | 106 |  | cluster5776 | 2036 | 109 |
| cluster13569 | 1457 | 109 |  | cluster5780 | 1544 | 113 |
| cluster13611 | 1929 | 97 |  | cluster5789 | 3690 | 100 |
| cluster13637 | 1572 | 109 |  | cluster5850 | 2748 | 109 |
| cluster13642 | 794 | 113 |  | cluster5895 | 3866 | 102 |
| cluster13651 | 1055 | 113 |  | cluster5918 | 2679 | 108 |
| cluster13675 | 1323 | 109 |  | cluster5927 | 1425 | 112 |
| cluster13704 | 1357 | 110 |  | cluster5935 | 2500 | 112 |
| cluster13724 | 2131 | 111 |  | cluster5949 | 1792 | 113 |
| cluster13733 | 1623 | 108 |  | cluster596 | 2888 | 109 |
| cluster1374 | 3103 | 89 |  | cluster5960 | 2339 | 108 |
| cluster13785 | 3150 | 103 |  | cluster5971 | 2374 | 109 |
| cluster13793 | 1556 | 107 |  | cluster5983 | 2271 | 109 |
| cluster13808 | 1759 | 108 |  | cluster6003 | 1642 | 111 |
| cluster13839 | 1880 | 103 |  | cluster6006 | 2256 | 113 |
| cluster13884 | 1197 | 113 |  | cluster6030 | 2235 | 109 |
| cluster13900 | 2326 | 109 |  | cluster6032 | 2497 | 110 |
| cluster13948 | 2837 | 100 |  | cluster6196 | 856 | 113 |
| cluster13967 | 750 | 112 |  | cluster6227 | 2563 | 109 |
| cluster13969 | 687 | 109 |  | cluster6244 | 4066 | 108 |
| cluster13980 | 479 | 113 |  | cluster6253 | 1465 | 113 |
| cluster13987 | 1018 | 113 |  | cluster6270 | 1619 | 112 |
| cluster14076 | 1960 | 106 |  | cluster6276 | 2382 | 109 |
| cluster14082 | 1525 | 109 |  | cluster6301 | 2785 | 108 |
| cluster14086 | 272 | 109 |  | cluster6310 | 2363 | 111 |
| cluster14124 | 1857 | 108 |  | cluster6328 | 2529 | 110 |
| cluster14129 | 946 | 111 |  | cluster6390 | 2577 | 110 |
| cluster14130 | 1114 | 111 |  | cluster6426 | 885 | 113 |
| cluster14149 | 1997 | 105 |  | cluster6434 | 1462 | 113 |
| cluster14175 | 822 | 112 |  | cluster6492 | 2498 | 111 |
| cluster14222 | 3726 | 102 |  | cluster6497 | 791 | 112 |
| cluster14228 | 2047 | 103 |  | cluster6581 | 1514 | 113 |
| cluster14236 | 842 | 113 |  | cluster6632 | 1579 | 113 |
| cluster14251 | 1025 | 113 |  | cluster6679 | 2153 | 110 |
| cluster14272 | 3040 | 97 |  | cluster6716 | 2119 | 109 |
| cluster14273 | 1884 | 102 |  | cluster6738 | 1557 | 113 |
| cluster14338 | 1467 | 111 |  | cluster6746 | 1574 | 110 |
| cluster14368 | 1466 | 107 |  | cluster6749 | 1414 | 113 |
| cluster14383 | 1481 | 111 |  | cluster6767 | 1522 | 113 |
| cluster14397 | 1689 | 107 |  | cluster6773A | 2479 | 110 |
| cluster14454 | 2003 | 106 |  | cluster6773B | 2257 | 109 |
| cluster14488 | 1629 | 109 |  | cluster6788 | 1298 | 112 |
| cluster14519 | 1935 | 104 |  | cluster6790 | 1328 | 113 |
| cluster14522 | 947 | 113 |  | cluster6795 | 2397 | 107 |
| cluster14525 | 504 | 113 |  | cluster6798 | 1537 | 113 |
| cluster14529 | 2568 | 107 |  | cluster6819 | 1295 | 113 |
| cluster14576 | 636 | 113 |  | cluster6837 | 2322 | 110 |
| cluster14624 | 2386 | 105 |  | cluster6864 | 1430 | 111 |
| cluster14629 | 2694 | 101 |  | cluster6883 | 2266 | 107 |
| cluster14676 | 843 | 113 |  | cluster6903 | 1612 | 111 |
| cluster14686 | 1746 | 108 |  | cluster6909 | 2226 | 112 |
| cluster14704 | 981 | 112 |  | cluster6910 | 1010 | 113 |
| cluster14739 | 1981 | 105 |  | cluster6924 | 1427 | 113 |
| cluster14778 | 2538 | 107 |  | cluster6937 | 1508 | 113 |
| cluster14781 | 920 | 113 |  | cluster6957 | 2164 | 112 |
| cluster14789 | 1640 | 107 |  | cluster6977 | 1329 | 113 |
| cluster14869 | 968 | 113 |  | cluster7009 | 1251 | 112 |
| cluster14882 | 1375 | 107 |  | cluster7016 | 2138 | 108 |
| cluster14913 | 1489 | 109 |  | cluster7043 | 1603 | 113 |
| cluster14916 | 1677 | 99 |  | cluster7062 | 2583 | 105 |
| cluster14924 | 1859 | 111 |  | cluster7075 | 2486 | 109 |
| cluster14930 | 1024 | 113 |  | cluster7092 | 1915 | 111 |
| cluster14934 | 607 | 112 |  | cluster7134 | 1990 | 110 |
| cluster14961 | 1499 | 112 |  | cluster7192 | 1421 | 112 |
| cluster14985 | 729 | 113 |  | cluster7198 | 2573 | 109 |
| cluster15015 | 612 | 111 |  | cluster7225 | 2040 | 113 |
| cluster15044 | 1747 | 107 |  | cluster7273 | 2534 | 109 |
| cluster15086 | 1781 | 108 |  | cluster7282 | 3585 | 108 |
| cluster15124 | 2201 | 107 |  | cluster7329 | 2041 | 109 |
| cluster15127 | 1764 | 108 |  | cluster7381 | 2237 | 105 |
| cluster15130 | 1562 | 110 |  | cluster7399 | 1050 | 113 |
| cluster15168 | 1430 | 108 |  | cluster7432 | 2989 | 110 |
| cluster15179 | 2360 | 109 |  | cluster7470 | 1924 | 109 |
| cluster15183 | 1467 | 108 |  | cluster7500 | 1966 | 111 |
| cluster15198 | 1670 | 106 |  | cluster7513 | 1469 | 112 |
| cluster15268 | 1746 | 107 |  | cluster7530 | 1504 | 110 |
| cluster15293 | 1364 | 105 |  | cluster7560 | 2221 | 108 |
| cluster15328 | 1905 | 110 |  | cluster7576 | 2253 | 111 |
| cluster15341 | 806 | 111 |  | cluster7587 | 1316 | 110 |
| cluster15383 | 1498 | 107 |  | cluster7598 | 3589 | 104 |
| cluster15387 | 1807 | 111 |  | cluster7616 | 1425 | 112 |
| cluster1543 | 1585 | 111 |  | cluster7621 | 3289 | 16 |
| cluster15467 | 1787 | 103 |  | cluster7627 | 1352 | 112 |
| cluster15565 | 1328 | 112 |  | cluster7651 | 1483 | 111 |
| cluster15567 | 705 | 113 |  | cluster7652 | 2092 | 109 |
| cluster15604 | 904 | 112 |  | cluster7662 | 726 | 111 |
| cluster15612 | 1998 | 106 |  | cluster7678 | 2670 | 109 |
| cluster15620 | 1303 | 111 |  | cluster7683 | 5133 | 103 |
| cluster15665 | 3300 | 59 |  | cluster7686 | 2141 | 113 |
| cluster15682 | 845 | 110 |  | cluster7707 | 1246 | 113 |
| cluster15724 | 1614 | 107 |  | cluster7712 | 1444 | 113 |
| cluster15732 | 1375 | 107 |  | cluster7718 | 1557 | 112 |
| cluster15830 | 2823 | 96 |  | cluster7722 | 1407 | 111 |
| cluster15869 | 801 | 112 |  | cluster7723 | 1562 | 112 |
| cluster15885 | 3346 | 96 |  | cluster7754 | 1569 | 112 |
| cluster15897 | 1641 | 112 |  | cluster7768 | 2607 | 105 |
| cluster15934 | 2059 | 107 |  | cluster7785 | 1988 | 111 |
| cluster15941 | 634 | 113 |  | cluster7819 | 1449 | 111 |
| cluster15951 | 1805 | 104 |  | cluster7821 | 727 | 94 |
| cluster15955 | 2626 | 109 |  | cluster7834 | 2323 | 109 |
| cluster15956 | 2695 | 107 |  | cluster7838 | 1239 | 112 |
| cluster15969 | 2034 | 105 |  | cluster7849 | 2275 | 107 |
| cluster15987 | 3149 | 102 |  | cluster7851 | 3446 | 103 |
| cluster15990 | 1634 | 106 |  | cluster7871 | 4609 | 94 |
| cluster16023 | 2121 | 103 |  | cluster7891 | 2126 | 110 |
| cluster16067 | 1453 | 108 |  | cluster7897 | 847 | 111 |
| cluster16073 | 3518 | 91 |  | cluster7898 | 1572 | 113 |
| cluster16107 | 1264 | 112 |  | cluster7919 | 804 | 113 |
| cluster16126 | 383 | 113 |  | cluster7922 | 1035 | 107 |
| cluster16128 | 730 | 113 |  | cluster7938 | 4074 | 105 |
| cluster16137 | 1338 | 108 |  | cluster7943 | 1502 | 113 |
| cluster16212 | 2718 | 107 |  | cluster7949 | 2191 | 110 |
| cluster16214 | 1577 | 97 |  | cluster7952 | 1299 | 113 |
| cluster16234 | 1623 | 104 |  | cluster7977 | 2514 | 110 |
| cluster16261 | 1950 | 108 |  | cluster8029 | 1382 | 112 |
| cluster16296 | 1310 | 110 |  | cluster8050 | 2387 | 110 |
| cluster16300 | 3448 | 82 |  | cluster8052 | 1868 | 107 |
| cluster16342 | 690 | 112 |  | cluster8081 | 1170 | 112 |
| cluster16356 | 1484 | 107 |  | cluster8100 | 4016 | 107 |
| cluster16370 | 812 | 112 |  | cluster8104 | 2421 | 106 |
| cluster16374 | 1938 | 105 |  | cluster8106 | 3061 | 96 |
| cluster16380 | 1179 | 111 |  | cluster8108 | 1388 | 111 |
| cluster16426 | 1693 | 108 |  | cluster8117 | 1404 | 113 |
| cluster16440 | 1845 | 102 |  | cluster8130 | 1484 | 112 |
| cluster16448 | 918 | 113 |  | cluster8176 | 2106 | 110 |
| cluster16500 | 1825 | 106 |  | cluster8210 | 4913 | 105 |
| cluster16509 | 891 | 113 |  | cluster8325 | 2023 | 113 |
| cluster16605 | 1218 | 111 |  | cluster8327 | 2073 | 104 |
| cluster16609 | 1455 | 110 |  | cluster8382 | 1000 | 113 |
| cluster16611 | 1913 | 83 |  | cluster8396 | 2072 | 112 |
| cluster16619 | 1659 | 110 |  | cluster8405 | 1880 | 111 |
| cluster16815 | 1264 | 109 |  | cluster8410 | 1346 | 111 |
| cluster16821 | 1866 | 101 |  | cluster8415 | 2560 | 108 |
| cluster16846 | 1490 | 103 |  | cluster8435 | 2086 | 110 |
| cluster16928 | 1024 | 111 |  | cluster8441 | 1657 | 112 |
| cluster16929 | 742 | 113 |  | cluster8460 | 1324 | 113 |
| cluster17012 | 895 | 113 |  | cluster8507 | 1385 | 113 |
| cluster17044 | 1673 | 108 |  | cluster8509 | 3495 | 108 |
| cluster17048 | 1915 | 101 |  | cluster8542 | 1747 | 110 |
| cluster17109 | 2146 | 104 |  | cluster8562 | 1578 | 112 |
| cluster17138 | 1529 | 107 |  | cluster8565 | 2179 | 109 |
| cluster17153 | 1753 | 100 |  | cluster859 | 2334 | 111 |
| cluster17197 | 1329 | 107 |  | cluster8600 | 3902 | 110 |
| cluster17230 | 637 | 113 |  | cluster8614 | 1365 | 111 |
| cluster17231 | 1158 | 113 |  | cluster8636 | 2177 | 112 |
| cluster17265 | 1501 | 109 |  | cluster8651 | 1022 | 113 |
| cluster1735 | 1555 | 111 |  | cluster8657 | 4865 | 109 |
| cluster17379 | 1889 | 95 |  | cluster8663 | 1821 | 107 |
| cluster17427 | 1356 | 109 |  | cluster8727 | 1635 | 111 |
| cluster17429 | 909 | 113 |  | cluster8773 | 2045 | 108 |
| cluster17441 | 1123 | 111 |  | cluster8798 | 1234 | 112 |
| cluster17455 | 1963 | 103 |  | cluster8837 | 2373 | 103 |
| cluster17501 | 975 | 111 |  | cluster8856 | 1177 | 113 |
| cluster17512 | 1232 | 109 |  | cluster8857 | 2065 | 110 |
| cluster17521 | 1499 | 110 |  | cluster8858 | 3907 | 108 |
| cluster17654 | 1264 | 110 |  | cluster8866 | 1286 | 112 |
| cluster17659 | 1733 | 98 |  | cluster8868 | 2718 | 108 |
| cluster17678 | 842 | 113 |  | cluster8872 | 1953 | 103 |
| cluster17735 | 1588 | 104 |  | cluster8890 | 3717 | 106 |
| cluster17900 | 1745 | 107 |  | cluster8913 | 1315 | 113 |
| cluster17975 | 727 | 113 |  | cluster8985 | 2856 | 92 |
| cluster18088 | 1894 | 103 |  | cluster9001 | 1233 | 108 |
| cluster18123 | 1830 | 104 |  | cluster9004 | 1359 | 113 |
| cluster18125 | 1836 | 106 |  | cluster9015 | 1131 | 112 |
| cluster18181 | 1142 | 109 |  | cluster9057 | 2176 | 106 |
| cluster18188 | 1652 | 32 |  | cluster9066 | 1317 | 112 |
| cluster18252 | 1697 | 101 |  | cluster9067 | 727 | 110 |
| cluster18257 | 1411 | 107 |  | cluster9075 | 1239 | 113 |
| cluster18260 | 1113 | 113 |  | cluster9099 | 4378 | 108 |
| cluster18374 | 1911 | 108 |  | cluster9103 | 1955 | 109 |
| cluster1844 | 2171 | 111 |  | cluster9107 | 2127 | 113 |
| cluster1870 | 2248 | 112 |  | cluster9114 | 2465 | 105 |
| cluster2091 | 2690 | 111 |  | cluster9126 | 1152 | 110 |
| cluster2133 | 2650 | 112 |  | cluster9133 | 1165 | 113 |
| cluster2216 | 8642 | 104 |  | cluster9145 | 1010 | 112 |
| cluster2219 | 3439 | 109 |  | cluster9163 | 2140 | 106 |
| cluster2233 | 3174 | 111 |  | cluster9164 | 1411 | 113 |
| cluster2305 | 2917 | 111 |  | cluster9173 | 2273 | 108 |
| cluster2375 | 2164 | 113 |  | cluster9175 | 1403 | 112 |
| cluster2386 | 2958 | 108 |  | cluster9198 | 2418 | 107 |
| cluster2408 | 2206 | 111 |  | cluster9247 | 1762 | 112 |
| cluster2424 | 2394 | 112 |  | cluster9308 | 1200 | 110 |
| cluster2428 | 4448 | 109 |  | cluster9313 | 2292 | 108 |
| cluster2430 | 2138 | 111 |  | cluster9388 | 2158 | 107 |
| cluster2433 | 1946 | 110 |  | cluster940 | 3711 | 11 |
| cluster2434 | 2209 | 113 |  | cluster9414 | 2071 | 109 |
| cluster2442 | 3078 | 112 |  | cluster9415 | 4188 | 111 |
| cluster2449 | 2249 | 107 |  | cluster9429 | 2017 | 108 |
| cluster2494 | 3598 | 113 |  | cluster9453 | 2708 | 105 |
| cluster2502 | 2240 | 113 |  | cluster9457 | 2166 | 107 |
| cluster2511 | 3858 | 105 |  | cluster9468 | 675 | 113 |
| cluster2517 | 2086 | 109 |  | cluster9510 | 3772 | 106 |
| cluster2534 | 1954 | 113 |  | cluster9514 | 2136 | 109 |
| cluster2555 | 3018 | 111 |  | cluster9543 | 1597 | 112 |
| cluster2558 | 4566 | 111 |  | cluster9559 | 1570 | 113 |
| cluster2573 | 2251 | 111 |  | cluster9591 | 1294 | 113 |
| cluster2588 | 2081 | 113 |  | cluster9595 | 1010 | 113 |
| cluster2597 | 2340 | 111 |  | cluster9600 | 3372 | 75 |
| cluster2601 | 1969 | 113 |  | cluster9622 | 5285 | 71 |
| cluster2615 | 2299 | 111 |  | cluster9667 | 3387 | 109 |
| cluster2627 | 2247 | 113 |  | cluster9677 | 1147 | 113 |
| cluster2647 | 3128 | 111 |  | cluster9686 | 3679 | 106 |
| cluster2648 | 3485 | 108 |  | cluster9687 | 2086 | 105 |
| cluster2649 | 2119 | 113 |  | cluster9689 | 1298 | 113 |
| cluster265 | 1266 | 102 |  | cluster9714 | 3985 | 108 |
| cluster2692 | 2301 | 111 |  | cluster9719 | 1061 | 113 |
| cluster2741 | 2237 | 113 |  | cluster9726 | 1980 | 110 |
| cluster2750 | 2146 | 112 |  | cluster9731 | 2028 | 108 |
| cluster2778 | 2033 | 113 |  | cluster975 | 4824 | 92 |
| cluster2784 | 2194 | 111 |  | cluster9753 | 1287 | 109 |
| cluster2807 | 1941 | 113 |  | cluster9781 | 716 | 113 |
| cluster2870 | 1640 | 113 |  | cluster9814 | 2048 | 108 |
| cluster2891 | 2927 | 109 |  | cluster9831 | 2100 | 109 |
| cluster2897 | 3446 | 109 |  | cluster9842 | 1690 | 113 |
| cluster2903 | 2749 | 113 |  | cluster9854 | 1704 | 111 |
| cluster2906 | 2349 | 112 |  | cluster9894 | 970 | 113 |
| cluster2915 | 3092 | 111 |  | cluster9906 | 1097 | 111 |
| cluster2937 | 2288 | 110 |  | cluster9926 | 1118 | 113 |
| cluster2960 | 2177 | 111 |  | cluster9937 | 2327 | 109 |
| cluster2962 | 2188 | 113 |  | cluster9959 | 1931 | 111 |
| cluster2968 | 1339 | 113 |  | cluster9967 | 3934 | 89 |
| cluster2981 | 2107 | 113 |  | cluster9995 | 2134 | 109 |

**Sup. Table 7.** Genes used for the time-calibrated phylogeny with their alignment length and number of sequences

| **Gene** | **length** | **# taxa** |
| --- | --- | --- |
| cluster11757 | 221 | 69 |
| cluster12457 | 351 | 67 |
| cluster12573 | 643 | 69 |
| cluster13724 | 177 | 68 |
| cluster13987 | 656 | 69 |
| cluster15387 | 689 | 69 |
| cluster2434 | 1842 | 69 |
| cluster2555 | 1146 | 67 |
| cluster2649 | 885 | 69 |
| cluster2784 | 1359 | 68 |
| cluster2807 | 121 | 69 |
| cluster3407 | 808 | 69 |
| cluster3680 | 548 | 69 |
| cluster3811 | 1279 | 69 |
| cluster3939 | 913 | 69 |
| cluster4129 | 895 | 68 |
| cluster4146 | 541 | 68 |
| cluster4298 | 798 | 69 |
| cluster5232 | 969 | 68 |
| cluster5440 | 1136 | 69 |
| cluster5780 | 680 | 69 |
| cluster6003 | 754 | 68 |
| cluster6032 | 1187 | 68 |
| cluster6328 | 1268 | 67 |
| cluster7723 | 597 | 68 |
| cluster7943 | 1090 | 69 |
| cluster8176 | 684 | 69 |

**Supplementary figures:**

**Fig. S1.** Testing for species richness differences across regions while accounting for differences in sampling intensity. A) Map of Neotropics indicating lowland and mountainous regions as defined by terrain ruggedness index (TRI). B) Rarefaction curves for estimated species richness showing differences in sampling effort in mountainous (N=3477) and lowland (N=1140) regions. C) Boxplot of rarefied species richness in lowland and mountainous regions, achieved by randomly drawing an equal number of occurrences from the two samples (N=1000), determining richness, and repeating this 1000 times.

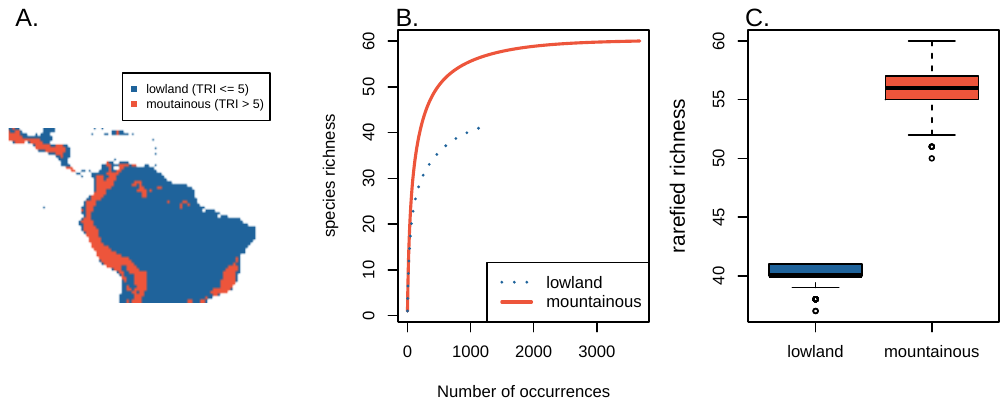

**Fig. S2.** Coalescent-based phylogeny. Numbers at branches indicate posterior probability.

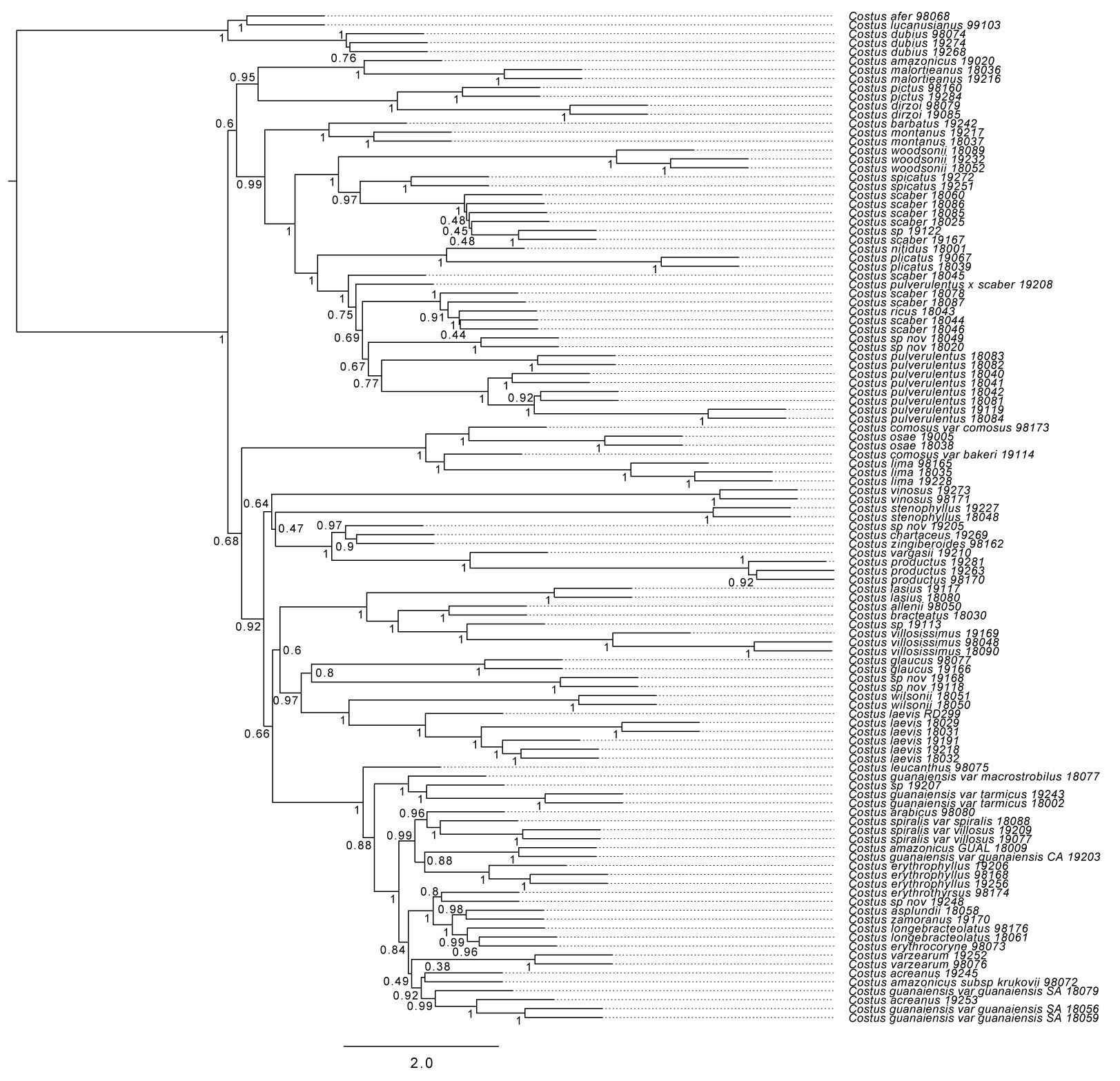

**Fig S3.** Agreement, conflict, and lack of signal of the individual 756 genes in relation with the concatenated topology. Colors in the pie charts indicate the number of genes that agree (blue), support a main alternative (green), conflict (red), or have no information in relation to the node in the concatenated tree.

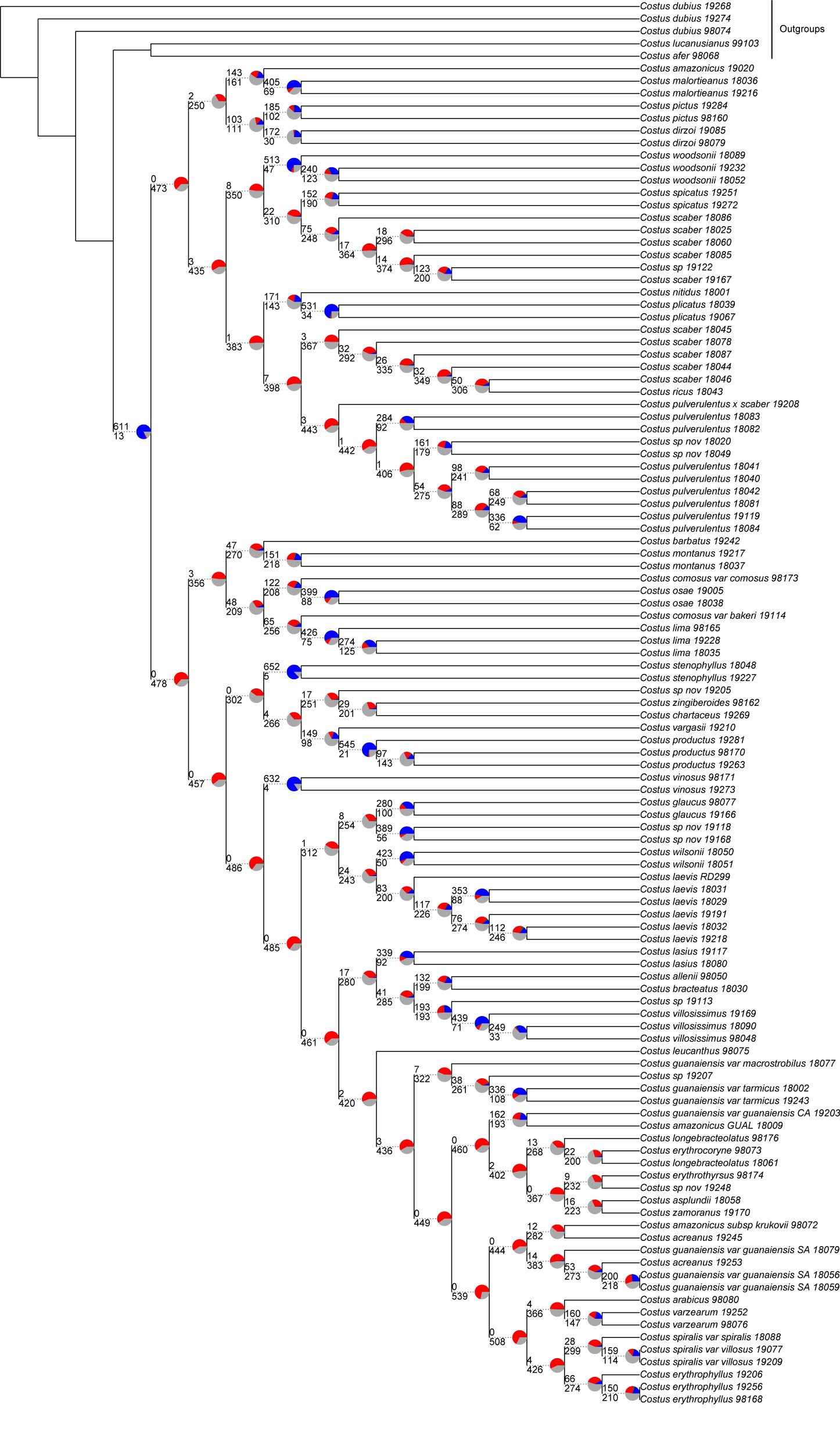

**Fig S4.** DEC biogeographic reconstruction of Neotropical *Costus*. Pie charts represent percent probabilities of areas for the ranges of ancestors. C: Central America, W: west indies, A: Andes, M: Amazon.

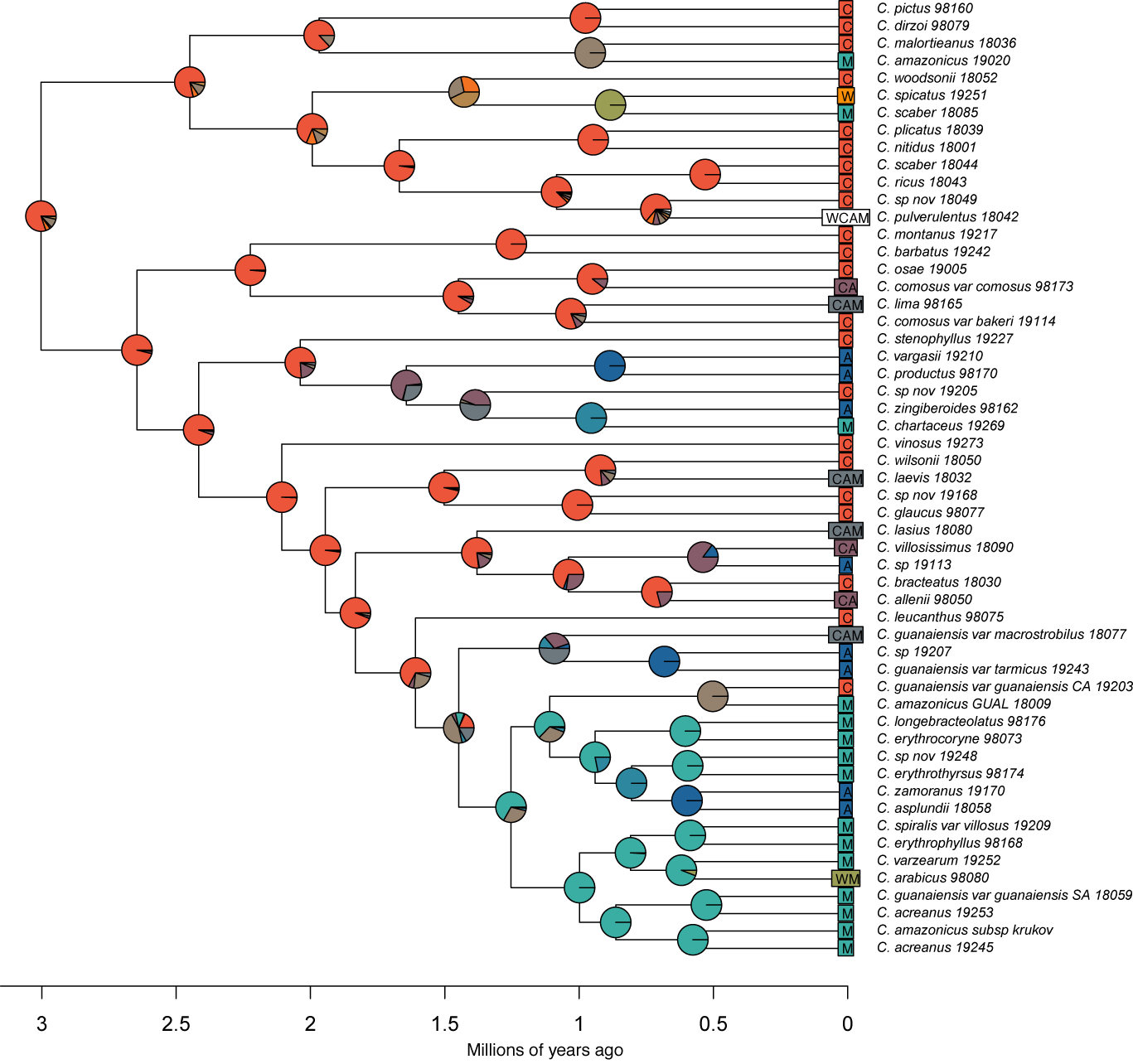

**Fig S5.** BAYAREALIKE biogeographic reconstruction of Neotropical *Costus*. Pie charts represent percent probabilities of areas for the ranges of ancestors. C: Central America, W: west indies, A: Andes, M: Amazon.

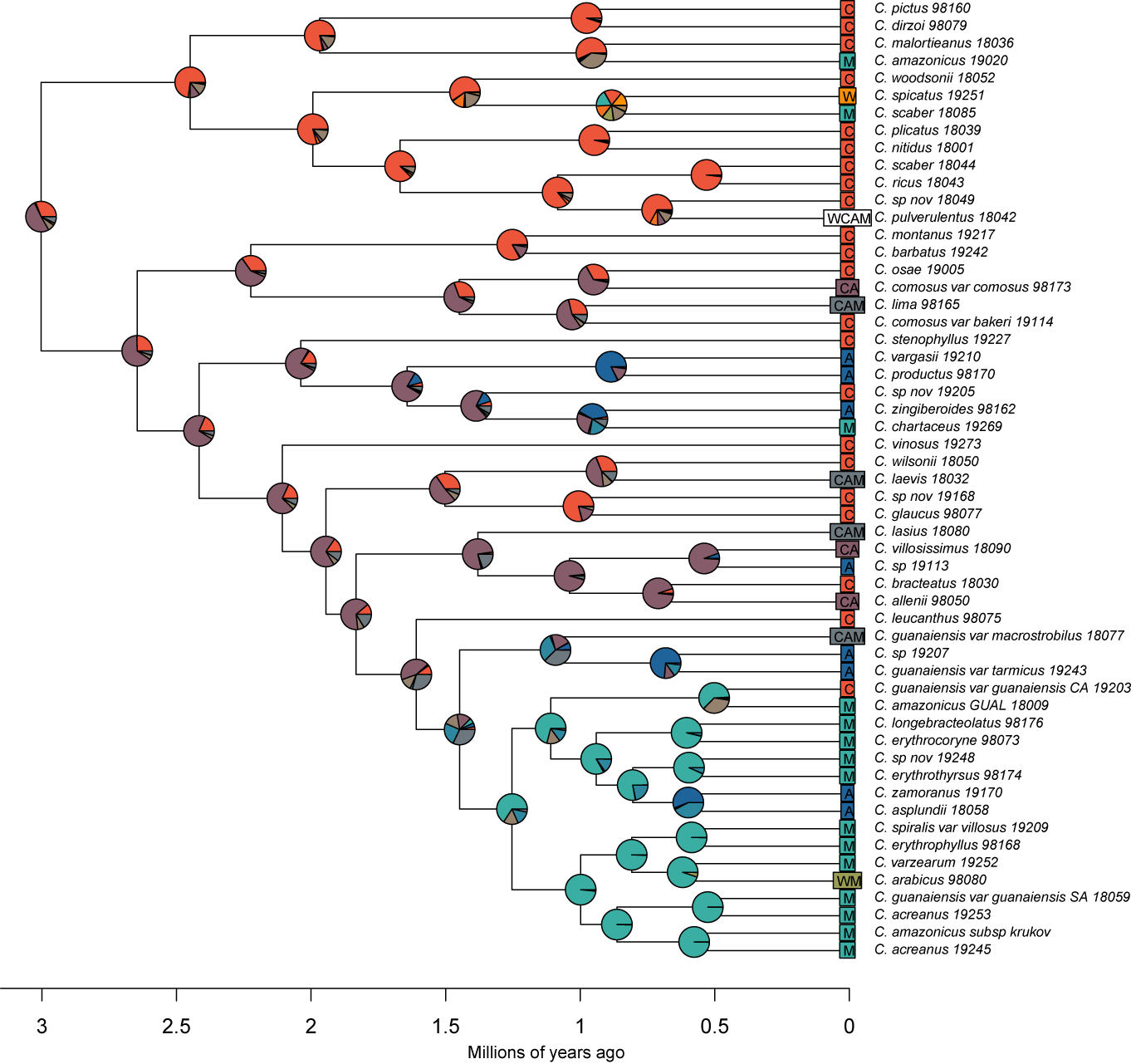

**Fig S6.** A) Bee and hummingbird pollination in extant species of *Costus*. Proportions of pollination syndromes are not significantly different between Amazonia and mountain-influenced species, *X^2^* (1 , N=55) = 1.51, *P* = 0.220. B) Pollinator transitions across all branches on the tree. Each branch is categorized as Amazonian or mountain-influenced according to the biogeographic reconstruction (see main text). There are no significant differences in the frequency of transitions to hummingbird pollination between Amazonian and mountain-influenced branches *Χ^2^* (NA , N=109) = 0.02, *P* = 1.

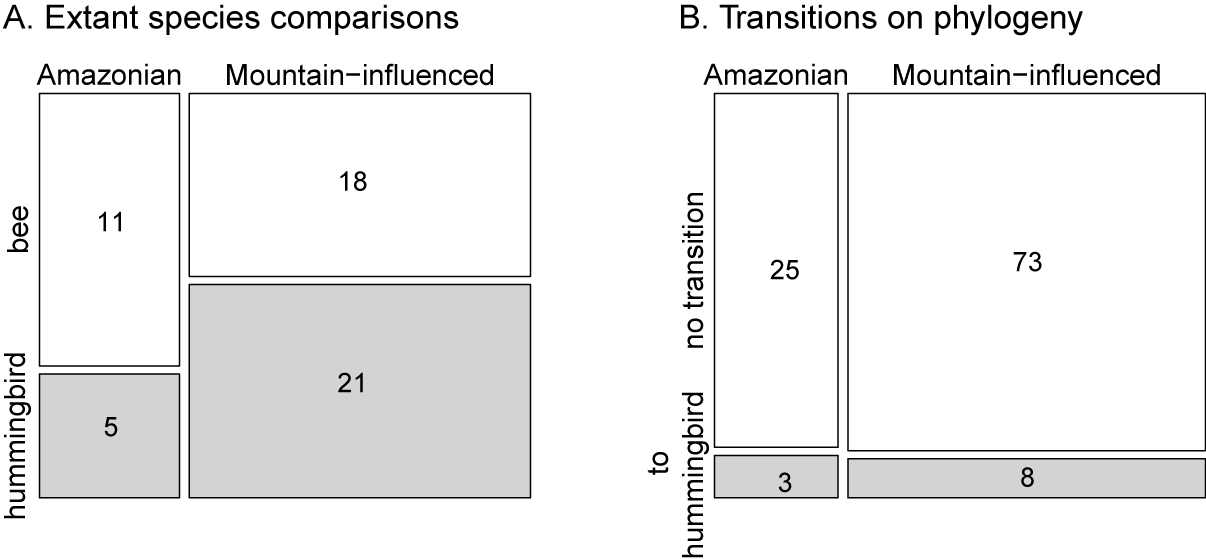

**Fig S7.** Distribution of sister species of *Costus.*

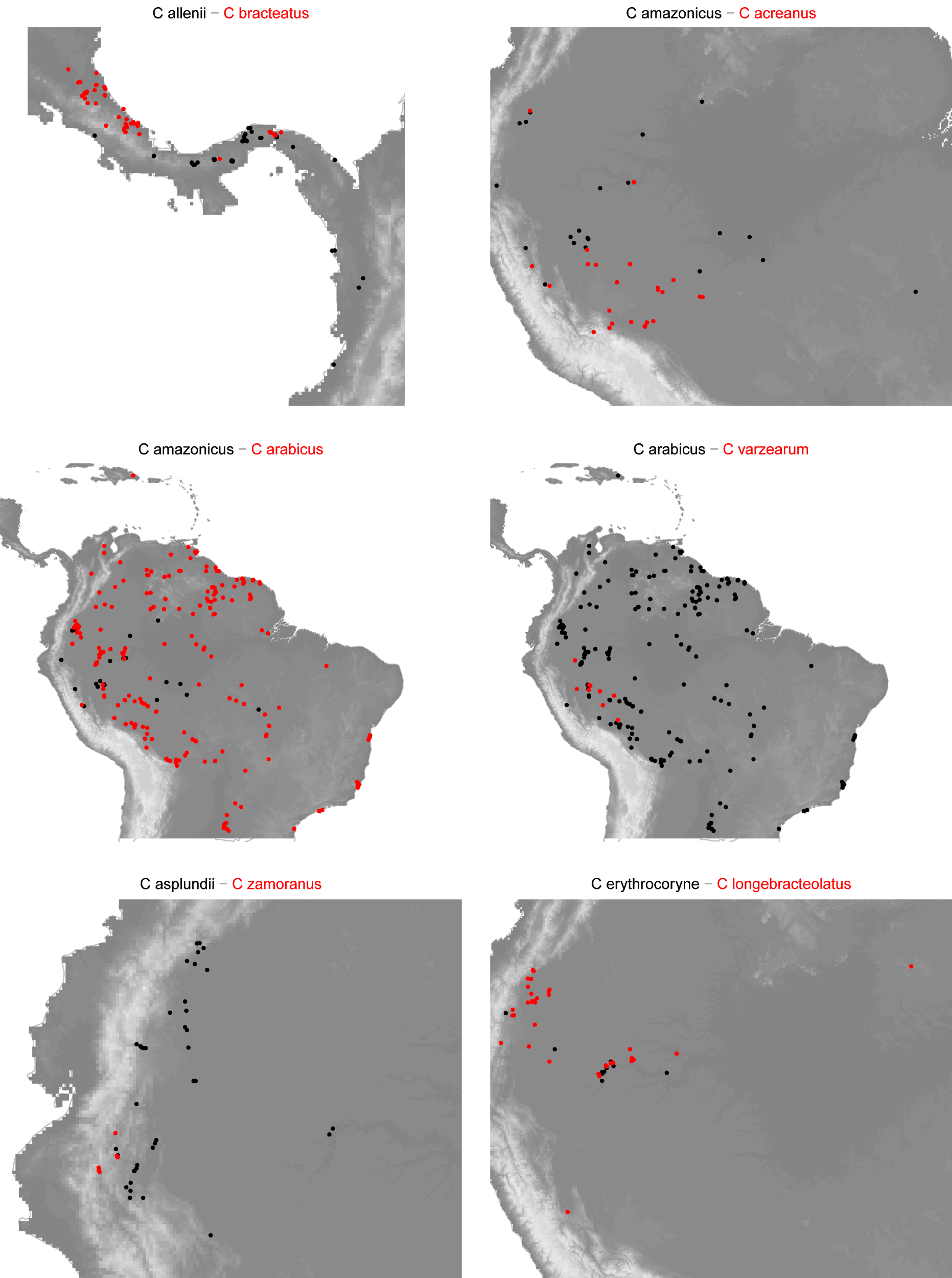

**Fig S7.** Distribution of sister species of *Costus* (continuation)*.*

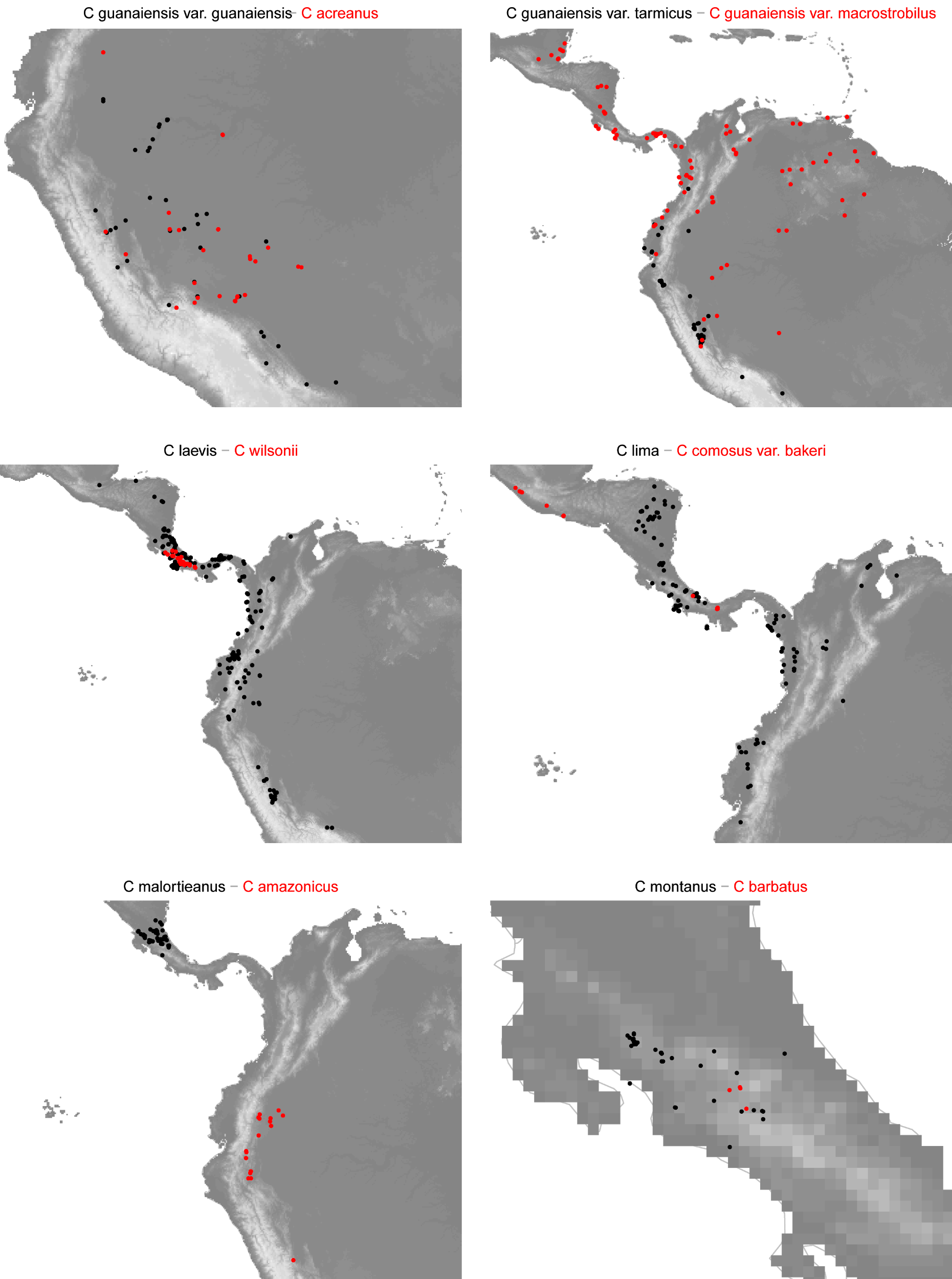

**Fig S7.** Distribution of sister species of *Costus* (continuation)*.*

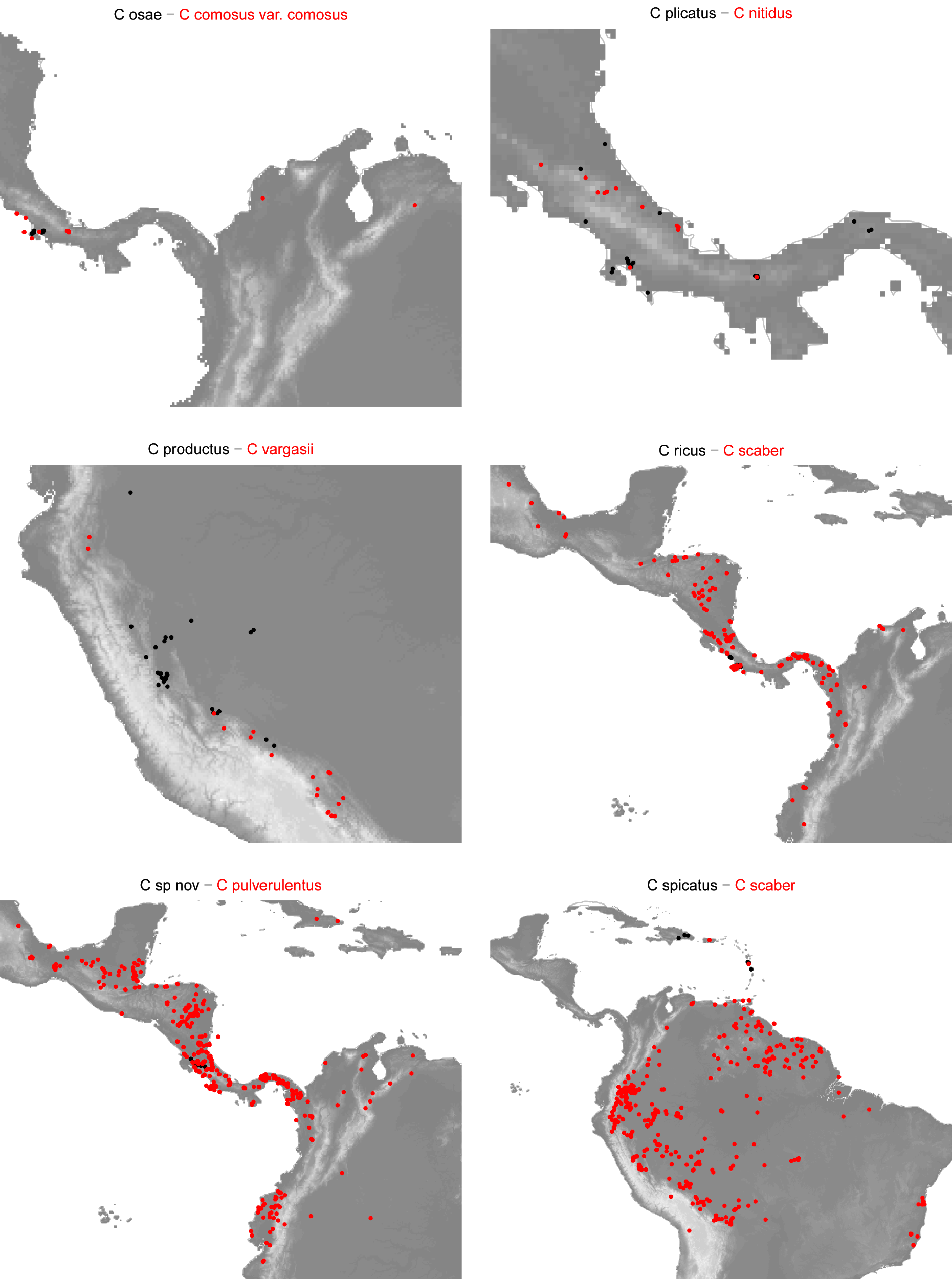

**Fig S7.** Distribution of sister species of *Costus* (continuation)*.*

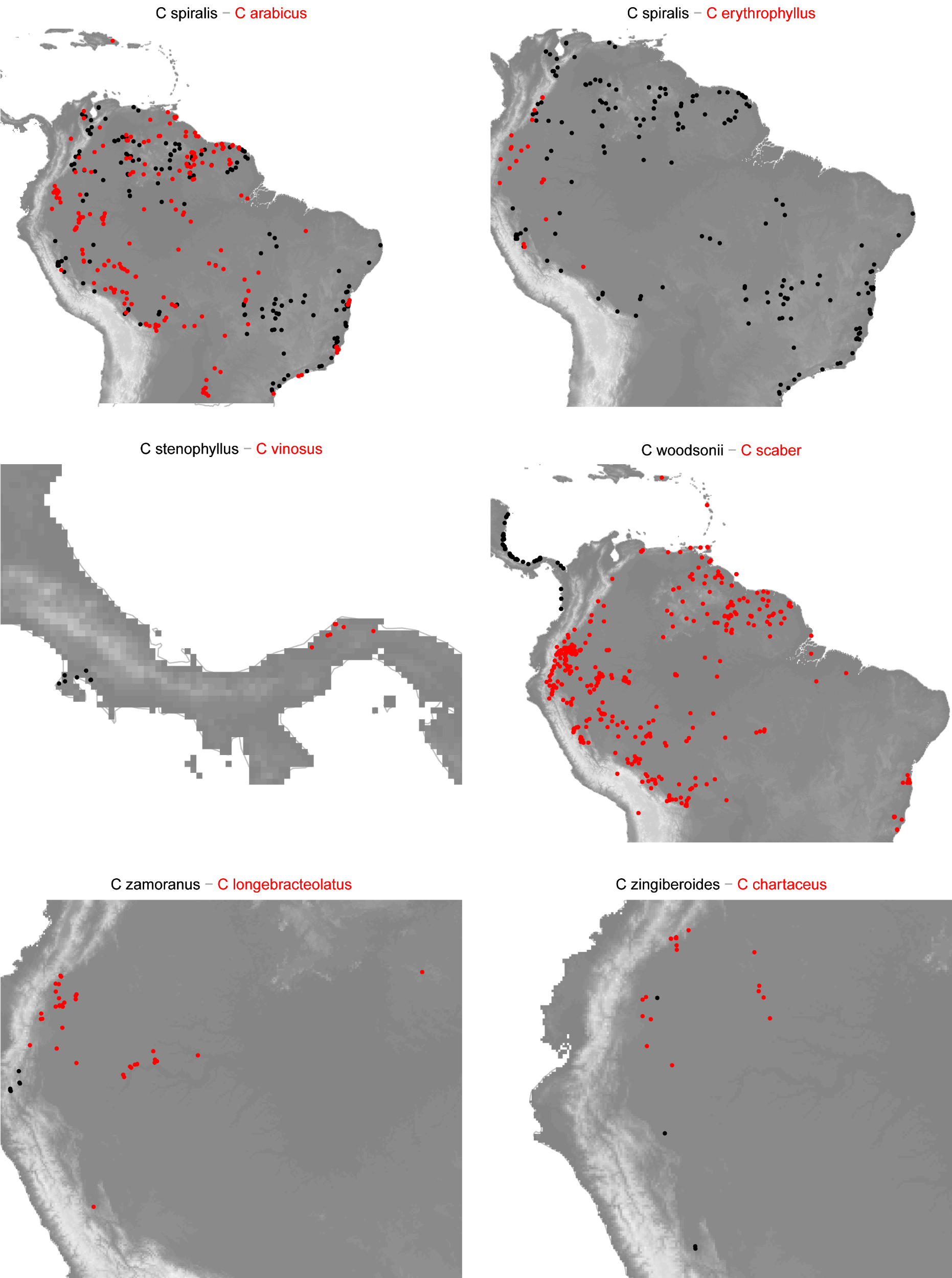

**Fig S8.** Niche comparisons of climatic niches in sister species of *Costus.* PC axes correspond to Fig. 5. Green lines show the distance between the centroids.

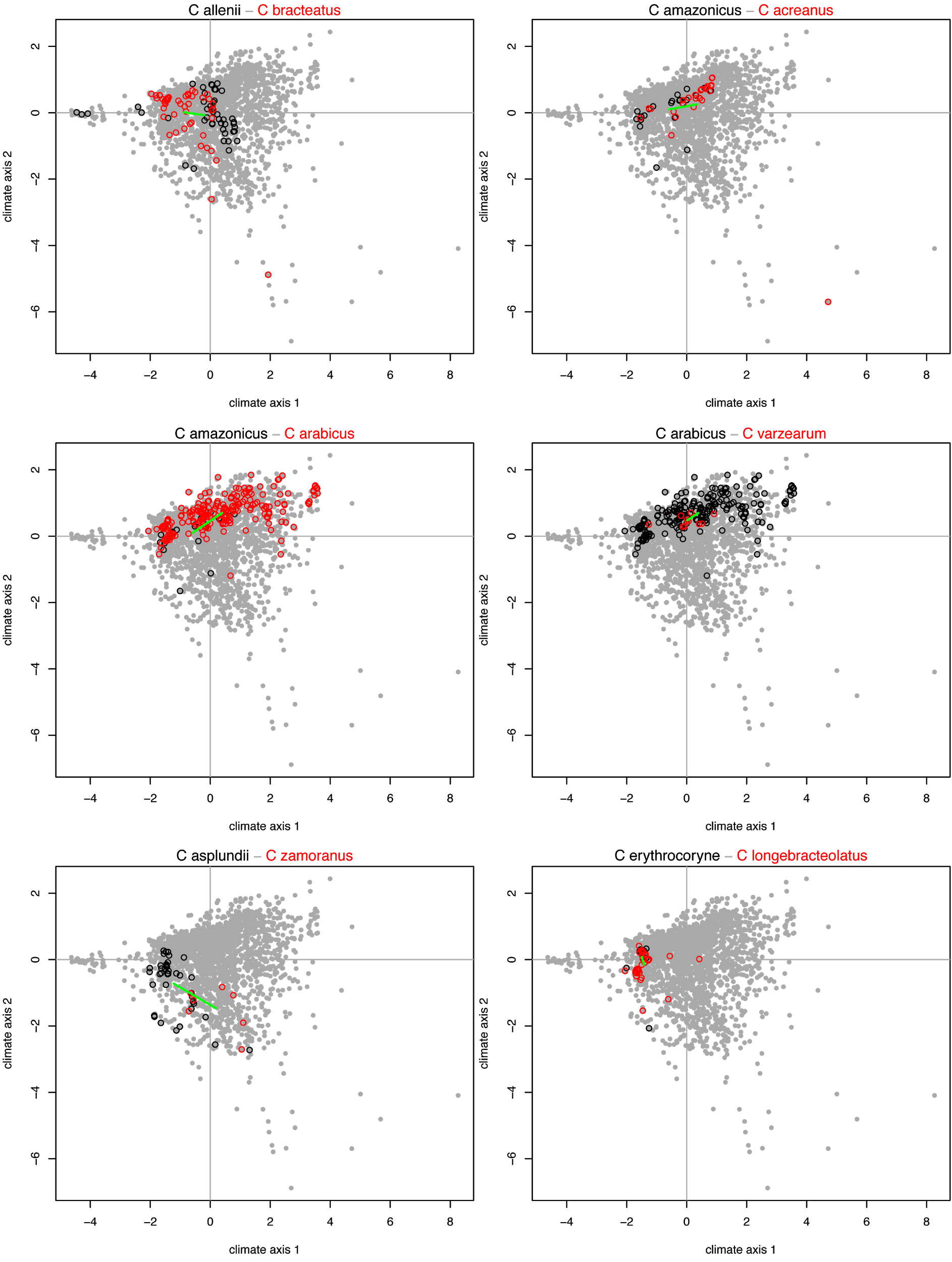

**Fig S8.** Niche comparisons of climatic niches in sister species of *Costus* (continuation)*.* PC axes correspond to Fig. 5. Green lines show the distance between the centroids.

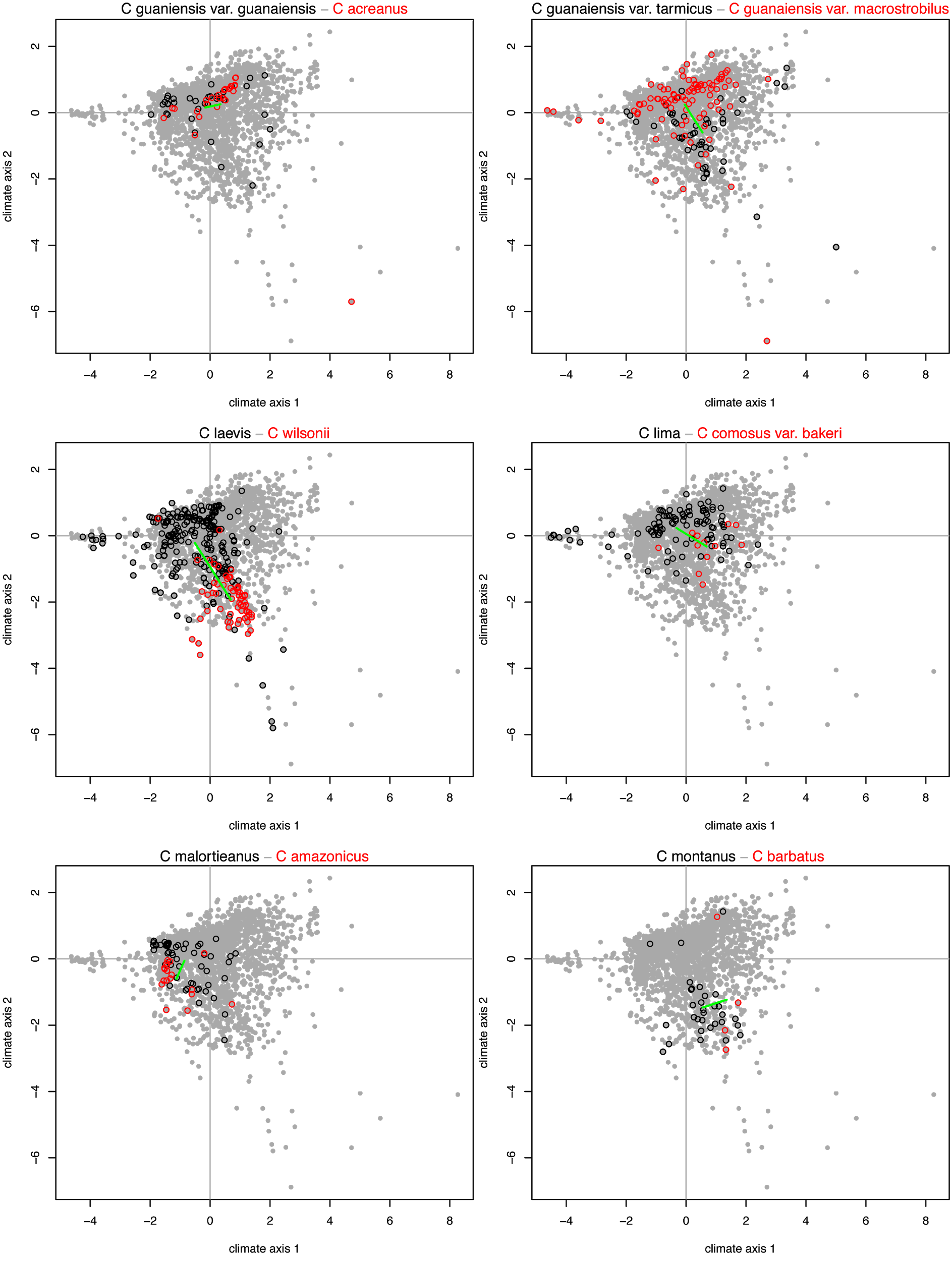

**Fig S8.** Niche comparisons of climatic niches in sister species of *Costus* (continuation)*.* PC axes correspond to Fig. 5. Green lines show the distance between the centroids.

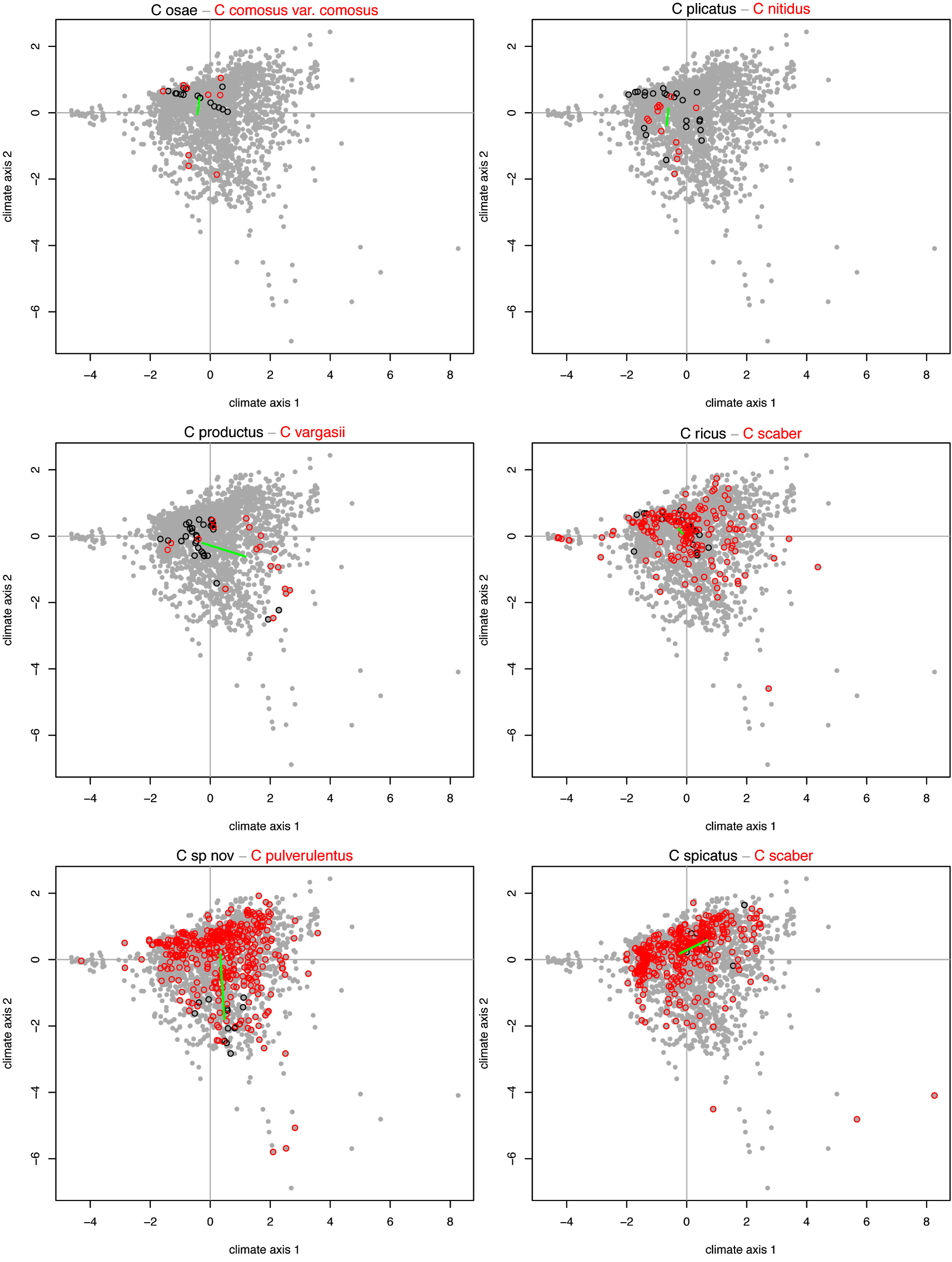

**Fig S8.** Niche comparisons of climatic niches in sister species of *Costus* (continuation)*.* PC axes correspond to Fig. 5. Green lines show the distance between the centroids.

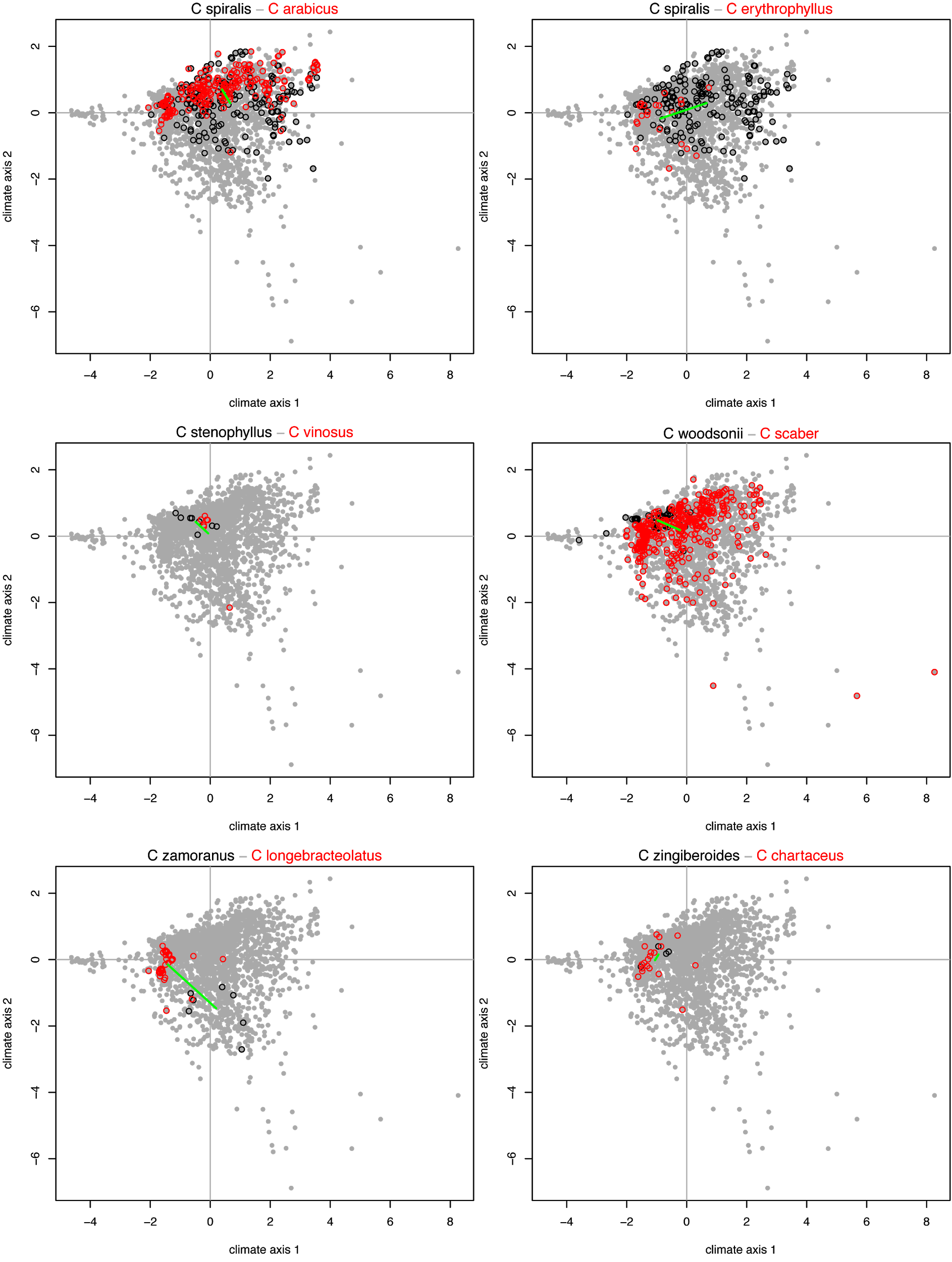

**Fig S9.** A) Comparison between the relative divergence time of mountain-influenced and Amazonian species pairs (two sample weighted T-test, t = -2.57, df = 20.7, *P* = 0.018); B) Range overlap v. divergence time (Mya) by region with marginal boxplots indicating differences in range overlap by region. C) Range size asymmetry v. divergence time (Mya) by region, with marginal boxplots indicating differences in range size asymmetry by region. In contrast with Fig 5 in main text, here range overlap and asymmetry were calculated on a grid of 0.1 decimal degrees.

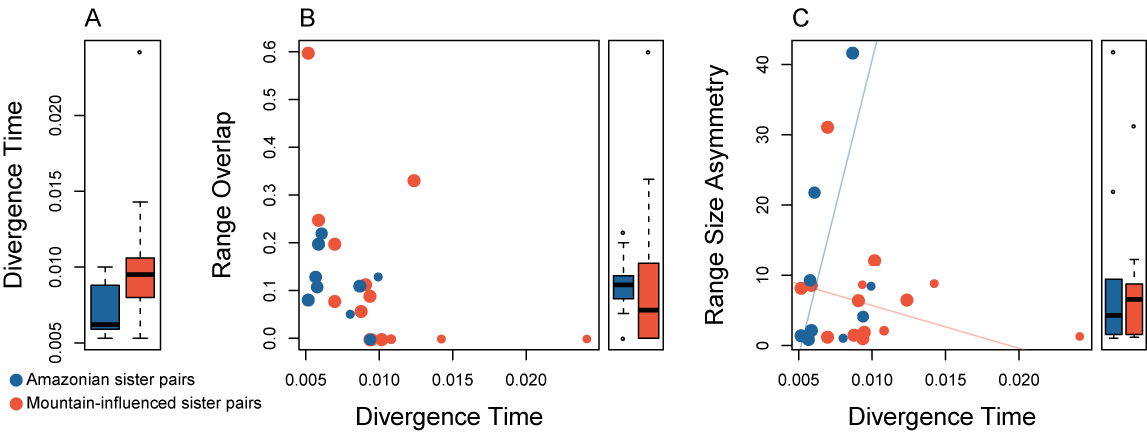
